# Agreement between structure prediction tools does not indicate biological plausibility in cancer fusion proteins

**DOI:** 10.64898/2026.11.20.768091

**Authors:** Ciarán Mc Gettrick, Gregory L. Wheeler

## Abstract

While protein-folding software has made great strides in recent years, reliability is known to be poorer on novel structures and failure modes are not fully characterised. Cancer fusion proteins present a unique case of proteins absent from model training data but constructed from parents with known structures. This study tests four popular protein-folding tools on three categories of protein: control, cancer-associated, and uncharacterised fusion proteins. We assessed computational performance, accuracy, and distinct failure modes. For well-characterised proteins, all tools tested returned the correct structures in all but one case, with some tools returning an average of two distinct structural forms in the exception. For characterised cancer-associated proteins, most predictions were highly accurate; poorer predictions often still returned some consistent high-confidence domains but produced divergent results in other areas that were both variable between tools and at odds with experimental structures. For cancer fusion proteins having available parent-protein reference structures, results fell into two clear categories. Roughly half contained structures matching to the functional domains of their parent proteins at high confidence, in keeping with biological expectation for these proteins; the others contained little evidence of matching structures, independent of the tool used. Predictions for fusion proteins differed far more between tools than those for characterised proteins with a median inter-tool TM-score 0.272 versus 0.853 respectively. However, agreement between tools did not indicate a more biologically plausible prediction. For example, all tools predicted drastically different structures for fusion protein TMPRSS2-ERG, but all predictions still contained the core ERG functional domains; conversely, all tools produced highly consistent structures for DNAJB1-PRKACA, but none of the predictions contained the expected functional regions. We recommend that users of these tools use care and caution in the assessments of outputs and consider biological knowledge before blindly trusting results.

## Introduction

Proteins are the primary functional molecules of the cell, and their three-dimensional structure is a major determinant of their biological activity (Branden and Tooze, 2012). Understanding protein structure is central to interpreting gene function, characterising disease mechanisms and identifying potential therapeutic targets. For decades, experimental methods such as X-ray crystallography (**?**), nuclear magnetic resonance (NMR) spectroscopy (Wüthrich, 1986) and cryo-electron microscopy (cryo-EM) (**?**) have served as the gold standard for determining protein structures at atomic resolution. However, these methods are slow, expensive (Yang and Zhang, 2015), and not universally applicable. Many biologically important proteins, particularly those that are intrinsically disordered (Wright and Dyson, 2015), membrane-associated, or existing only transiently as part of larger complexes remain experimentally unresolved. As of 2024, the Protein Data Bank (PDB) contained experimentally determined structures for greater than 227,000 proteins (Burley et al., 2025), while simultaneously the number of protein sequences in UniProtKB exceeded 246 million (UniProt Consortium, 2025). This is more than a 1000-fold difference and it presents a significant challenge that experimental methods alone cannot realistically close.

Computational structure prediction has for a long time, been pursued as a means of bridging this gap. Early approaches relied on homology modelling, for which the structure of a target protein is inferred from sequence similarity to a protein with a known experimental structure (Martí-Renom et al., 2000). While effective for proteins with close homologues, homology modelling is fundamentally limited by the availability of suitable templates and performs poorly on novel or uncharacterised sequences (Baker and Sali, 2001). Threading and fragment assembly methods, such as Rosetta (Rohl et al., 2004), extended the reach of computational prediction beyond close homologues, but accuracy remained inconsistent, particularly for proteins with novel folds or limited evolutionary conservation. The Critical Assessment of Protein Structure Prediction (CASP) experiments, held biennially since 1994, have provided a standardised framework for evaluating progress in this field (Kryshtafovych et al., 2019).

The field was greatly transformed by the development of deep-learning approaches to structure prediction. The most prominent was AlphaFold2, which achieved near-experimental accuracy on a wide range of targets at CASP14 in 2020 (Jumper et al., 2021; Kryshtafovych et al., 2021). AlphaFold2 integrates a multiple sequence alignment (MSA) of evolutionary relatives with a neural network architecture, the Evoformer, to predict inter-residue relationships from which a three-dimensional structure is generated. Its success prompted the release of the AlphaFold Protein Structure Database. This now provides predicted structures for over 200 million protein sequences (**?**), effectively closing the numerical gap between known sequences and available structural models, at least in terms of coverage, if not guaranteed accuracy. Demis Hassabis and John Jumper were subsequently awarded half of the 2024 Nobel Prize in Chemistry for their contributions to computational protein structure prediction, with the other half being awarded to David Baker for computational protein design.

Following AlphaFold2, a number of alternative prediction tools have emerged, each making distinct architectural choices that affect their applicability and ability to different use cases. AlphaFold3, the successor to AlphaFold2, replaces the structure module with a diffusion-based architecture and extends prediction capabilities to protein–ligand, protein–nucleic acid and protein–protein complexes (Abramson et al., 2024). Boltz-1 adopted a similar diffusion-based approach while offering an open-source, locally deployable implementation (Wohlwend et al., 2024), and its successor Boltz-2 further extended this to include binding-affinity prediction (Passaro et al., 2025). In contrast, ESMFold (Lin et al., 2023) and OmegaFold (Wu et al., 2022) take a fundamentally different approach. Rather than relying on MSAs constructed from databases of evolutionary homologues, these tools derive structural features directly from protein language models trained on large sequence history (Rives et al., 2021). This enables single-sequence prediction without an external sequence-database search. This distinction is not just a matter of convenience; it has direct implications for the types of proteins these tools can meaningfully predict. MSA-dependent tools derive much of their predictive power from evolutionary conservation signals present across families of related sequences (Rao et al., 2021). Novel proteins with no known homologues may have little or no evolutionary information available. Mutated or fusion proteins may retain evolutionary information for their known regions, but not for altered residues or novel fusion junctions. The relative performance of MSA-dependent and MSA-free tools on these proteins is therefore not well understood.

A critical but often overlooked aspect of structure-prediction tool evaluation is the distinction between prediction accuracy and prediction reliability. Each of the tools discussed above produces internal confidence metrics alongside its structural predictions: predicted local distance difference test (pLDDT) scores estimate per-residue accuracy, while predicted TM-scores (pTM) and predicted aligned error (PAE) matrices provide global and pairwise confidence assessments, respectively (Jumper et al., 2021). These metrics are routinely used by researchers to judge whether a given prediction should be trusted. However, they are generated by the model itself without reference to experimental data, and their relationship to true structural accuracy is not guaranteed. A model that is consistently overconfident poses a greater practical risk than a model that is less accurate overall but honest about its own uncertainty, because the former may lead researchers to place unwarranted trust in incorrect structures. Despite its importance, systematic evaluation of confidence calibration—whether a tool’s self-reported confidence reliably predicts its actual accuracy—has received less attention than overall accuracy in the benchmarking literature, which has focused predominantly on accuracy rankings such as those provided by CASP (Kryshtafovych et al., 2021).

This gap is particularly relevant in the context of cancer genomics, where structure prediction is increasingly applied to proteins for which limited or no experimental structural data exist (Zhao et al., 2024). Cancer-associated proteins, including oncogenes, tumour suppressors and the products of cancer-specific gene fusions, are of direct therapeutic and diagnostic interest (Hanahan, 2022; Vogelstein et al., 2013). However, many are large, multi-domain signalling proteins whose structural complexity may pose challenges distinct from those of the small, well-characterised proteins typically used in benchmarking studies. Furthermore, cancer gene fusions—chimeric proteins formed by the aberrant joining of two normally separate genes (Mertens et al., 2015)—combine protein regions that evolved separately. Although the individual regions may have homologues, the fusion junction and overall domain arrangement may have no direct evolutionary precedent, creating particular challenges for structure-prediction tools (Kumar et al., 2024; **?**). The FusionPDB database has demonstrated the feasibility of large-scale structure prediction for human fusion proteins using AlphaFold2, providing thousands of predicted fusion-protein structures and identifying potential druggable targets through downstream structural and therapeutic analyses (Kumar et al., 2024). However, systematic comparison of multiple structure-prediction tools on fusion proteins remains limited.

To address this, this study compares four current AI-based protein structure prediction tools: AlphaFold3 (Abramson et al., 2024), Boltz-2 (Passaro et al., 2025), ESMFold (Lin et al., 2023) and OmegaFold (Wu et al., 2022). These tools are applied to three biologically motivated categories of human proteins: well-characterised structural benchmark proteins, which act as the control group, cancer-associated proteins and cancer-derived fusion proteins. Protein sequences were derived from the MANE RefSeq consensus transcript set (Morales et al., 2022), and prediction accuracy was evaluated against experimentally determined reference structures from the Protein Data Bank (Berman et al., 2000) using the US-align structural comparison framework (Zhang et al., 2022). Rather than focusing solely on prediction accuracy, this study takes confidence calibration as its primary analytical lens, asking not only which tool produces the most accurate structures, but which tool most reliably knows when its predictions can be trusted. Specifically, this study aims to (1) evaluate the confidence calibration of each tool by quantifying the relationship between self-reported confidence and true structural accuracy across protein categories, with particular attention to cancer-associated proteins, and (2) evaluate inter-tool structural agreement on fusion proteins for which no experimental ground truth exists, using consensus between independently developed prediction architectures as a proxy for prediction reliability.

## Methods

### Collection of reference sequence and structural data

A total of 60 proteins were included in this study. 20 characterised benchmark, control proteins, 20 cancer-associated proteins and 20 fusion proteins. The characterised control proteins were assembled during an initial exploratory phase and represents a range of well-studied proteins with varied lengths, functions and structural characteristics. As this set was assembled pragmatically rather than through systematic database sampling, it is treated as a curated benchmark panel.

The cancer-associated set was selected separately on the basis of established relevance to cancer biology, also on the likely availability of experimentally determined structures and computationally manageable sequence lengths. The selected proteins represented a range of oncogenes, tumour suppressors, receptors, signalling proteins and apoptosis regulators. Amino acid sequences for the cancer-associated proteins were extracted from the MANE GRCh38 v1.5 RefSeq protein dataset (Morales et al., 2022). These sequences were combined with the characterised control and fusion-protein sequences to create a fixed master FASTA manifest. Proteins 1 to 20 represented the characterised control set, proteins 21 until 40 represented the cancer-associated set and proteins 41–60 represented the cancer-derived fusion set. Each sequence was then treated as an independent monomer. Identical sequence inputs and identifiers were used across all four prediction workflows.

Experimentally determined reference structures were collected for the 40 non-fusion proteins. Initial candidate structures were identified by querying the PDBe best structures service using the corresponding protein accession. This service ranks structures deposited in the Protein Data Bank according to factors including sequence coverage and experimental resolution (Armstrong et al., 2020). The highest-ranked structure was used as an initial candidate rather than being accepted automatically.

Each candidate was manually validated at the level of individual chains. The amino acid sequence represented by the candidate PDB chain was extracted using Biopython (version 1.84) (Cock et al., 2009) and aligned against the complete sequence supplied to the prediction tools. A chain was accepted if it shared at least 95% sequence identity with the target and at least 95% of the chain aligned to the target sequence. Structures containing an incorrect chain, an incompatible isoform, low sequence identity or insufficient coverage of the target protein were rejected, and an alternative reference was selected when available. This validation was particularly important for proteins such as KRAS and BCL-xL, for which initial automated assignments identified inappropriate chains within multichain PDB entries.

Reference structures were also required to have been determined by X-ray crystallography or cryo-EM at a resolution of 4 Å or better. Structures were also required not to be complexes in which a bound partner holds the target in a non-native conformation. Engineered point mutations in the reference chain were allowed. Five structures from the initial search did not meet these criteria and were replaced with the next-ranked structure that did (Supplementary Table S1). Reference selection was therefore based only on the experimental method, resolution, sequence and binding state of each structure, and was performed independently of prediction performance. Results obtained with the originally retrieved structures are given in Supplementary Table S2.

The top-ranked calmodulin-1 structure (9MXD) is calmodulin bound to a myosin light-chain kinase peptide, and was replaced with calcium-bound calmodulin (1CLL). Because calmodulin changes shape substantially upon calcium binding, predictions were also compared with calcium-free calmodulin (1CFD), as described in the Results.

Reference structures were classified according to the percentage of the target sequence represented by the validated experimental chain.

References covering at least 90% of the target sequence were classified as near-full-length structures (n = 17) and were included in global structural-accuracy analyses. This threshold permitted the absence of short flexible loops or terminal regions while retaining coverage of the overall tertiary structure.

References covering 20–89% of the target sequence were classified as partial or domain-level structures (n = 21) and were retained for regional structural comparisons. Predictions were assessed only over the experimentally resolved region in these cases, preventing models from being penalised for residues that were absent from the reference structure.

References covering less than 20% of the target sequence were classified as short fragments and excluded from the primary structural-accuracy analysis because isolated motifs provide poor topological information for evaluating overall fold accuracy. Insulin and polyubiquitin-C met this exclusion criterion. Both proteins were retained in the computational-performance analysis, but were excluded from comparisons against experimental structures. This produced a final dataset of 38 proteins with validated experimental references. The method, resolution, engineered mutations and bound partners of each reference are listed in Supplementary Table S1.

### Inference of Protein Structure from Sequence Data

Protein structure predictions were generated using four computational workflows: ESMFold, Boltz-2, OmegaFold and AlphaFold3. All workflows were run on the University of Galway School of Mathematical and Statistical Sciences’ on-premises high-performance computing cluster, with jobs submitted and managed using the SLURM workload manager. The fixed master FASTA file was used as the sequence source for every workflow, and all 60 proteins were processed in a batch, with each protein predicted separately as an individual monomer.

#### AlphaFold3

AlphaFold3 version 3.1.0 was run using a Singularity container, built from the AlphaFold3 GitHub repository (commit f43a5c4, 17 July 2026) via the included Dockerfile. Pre-trained model weights were obtained from Google DeepMind via academic-use license (8 July 2026). Reference database files were obtained using the provided fetch databases.sh script on 19 July 2026.

Before running AlphaFold3, FAA-formatted sequence data was converted into individual-protein JSON input files using a custom Python script. AlphaFold3 (**?**) was then run as a two-phase process, using the pre-built container run via Singularity. Weights were obtained from Google DeepMind under its research use agreement. The first, longer-running phase was run on CPU (16 cores, 64 GB of memory) and mainly performed the multiple sequence alignment (MSA) search. The second phase performed structure inference on a single NVIDIA A100 40 GB GPU, additionally using 2 CPU cores and allocated 32 GB of CPU memory. Predictions were performed using a single model seed (42), and the top-ranked model for each protein was used.

#### Boltz-2

Boltz-2 version 2.2.1 (Passaro et al., 2025) was installed in a dedicated Conda environment and run on the CPU partition. The SLURM job allocated one node, 16 CPU cores and 96 GB of RAM. Each protein was submitted independently to the Boltz-2 model on CPU, using 16 processing threads, the integrated ColabFold MSA server (Mirdita et al., 2022) and random seed 42. All other settings, including the number of recycling and sampling steps, were left at the default. One independent monomeric structure prediction was created for each of the 60 proteins. Predicted structures and accompanying confidence outputs were retained for analysis. The extracted outputs included the overall confidence score, pTM, complex pLDDT and complex predicted distance error (pDE), together with protein length, execution device, runtime, system-memory use, CPU time, CPU-core use, output-file location and completion status.

#### ESMFold

ESMFold was run using fair-esm version 2.0.0 (Lin et al., 2023), installed in a dedicated Conda environment. The SLURM job was allocated one node, 32 CPU cores and 160 GB of RAM. The ESMFold v1 model was loaded with a chunk size of 64 and used to generate one predicted structure in PDB format for each of the 60 protein sequences using an automated batch workflow. The workflow also recorded the sequence length, execution device, completion status, wall-clock runtime, CPU time, number of CPU cores used and system-memory use after each prediction. Model confidence outputs, including mean predicted local distance difference test (pLDDT), predicted TM-score (pTM) and mean predicted aligned error (PAE), were extracted and written to a consolidated results file. Successful model generation was verified by confirming the presence of the expected PDB output for each protein.

#### OmegaFold

OmegaFold (Wu et al., 2022) was installed from its GitHub repository (commit 313c873, December 2022) in a dedicated Conda environment and run on the CPU partition with default settings. The SLURM job allocated ones node, 16 CPU cores and 128 GB of RAM. The same 60 amino acid sequences were run, with each prediction written to a separate protein-specific output directory. A single structural model was retained for each target. Per-residue confidence values encoded in the predicted structure files were extracted and averaged to obtain a mean pLDDT value for each protein. OmegaFold does not provide a directly comparable pTM output, so it was not included in the pTM-based confidence analysis. Runtime, resource use and completion status were recorded by the associated SLURM workflow.

### Performance benchmarking

#### Resource usage and runtime

Computational performance was monitored using custom Python workflows and the SLURM workload manager. Runtime was recorded separately for each protein. For ESMFold, CPU time (user plus system time) and memory use were recorded from the running Python process, with memory measured after each prediction rather than as peak use. For OmegaFold and Boltz-2, CPU time and peak RAM consumption were obtained using GNU time. Effective CPU-core use was then calculated as CPU time divided by runtime. Protein length, execution device and completion status were recorded alongside these measurements.

ESMFold, OmegaFold and Boltz-2 were evaluated using CPU inference, although their allocated resources differed: ESMFold was allocated 32 CPU cores and OmegaFold and Boltz-2 were allocated 16. CPU time was then therefore used as the main measure of computational cost. AlphaFold3 used a CPU-based data-processing stage followed by GPU inference and was therefore considered separately rather than treated as directly hardware equivalent. Resource measurements covered structure prediction only and therefore did not include subsequent US-align comparisons or statistical analysis.

#### Structural prediction accuracy

Predicted structures were compared with the experimental reference structures using US-align (version 20260527) (Zhang et al., 2022) with default settings. Before each comparison, the predicted model was trimmed to the target residues covered by the reference, and the reference chain was trimmed to its resolved residues, so that both structures represented the same region. The TM-score normalised by the length of the experimental structure was used as the main measure of accuracy. RMSD, aligned residue count and sequence identity were also recorded to provide additional information about each alignment.

Near-full-length references were used to assess overall fold accuracy, while partial references were used to assess only the experimentally resolved region. The analysis included 38 proteins. Insulin and polyubiquitin-C were excluded because their available experimental structures covered less than 20% of the supplied sequences.

#### Self-assessed confidence accuracy

Each tool’s reported confidence was then compared with its measured accuracy against the experimental structures. For ESMFold, Boltz-2 and AlphaFold3, the accuracy–confidence difference was calculated for each protein as the reference-normalised TM-score minus the model-reported pTM. A positive value indicates that the tool underestimated its own accuracy, and a negative value that it overestimated it. Custom Python scripts were used to match the confidence measurements with the corresponding US-align results, and are available through the project’s GitHub repository.

### Performance on novel chimeric structures vs. known proteins

The 20 fusion proteins were selected from FusionPDB2 https://pubmed.ncbi.to to represent recurrent cancer-associated fusions. Each fusion was predicted from amino acid sequence as a single monomer by all four tools using the same settings as the non-fusion proteins.

As experimental structures are not available for the fusion proteins, predictions were instead compared with experimental structures of the two parent proteins from which each fusion is derived. A reference structure was selected from the PDB for each parent, and the percentage of each parent’s sequence retained in the fusion was calculated. The retained proportions were sharply divided: most parents shared either 10–20% or more than 90% of their sequence with the fusion, with a single intermediate case at approximately 40%. Parents sharing less than 90% of their sequence were excluded, as a score based on a small region would not represent accuracy across the fusion. This left 12 parent regions from 11 fusions (6 Parent A and 6 Parent B), with RUNX1-RUNX1T1 the only fusion for which both parents were retained. Each predicted fusion was aligned to each retained parent structure using US-align, and the TM-score normalised by the length of the parent structure was used as the measure of accuracy.

To measure agreement between tools, the four predicted structures for each protein were compared with one another in every pairwise combination using US-align, for both the 20 fusion proteins and the 40 non-fusion proteins. Each pair was compared in both directions, and all pairwise TM-scores for a protein were averaged to give a single inter-tool similarity score, so that each protein contributed one value to the statistical analysis. As this comparison does not require an experimental reference, insulin and polyubiquitin-C were included.

### Statistical analysis

Statistical analyses were carried out in GraphPad Prism (version 11.1.0). As TM-scores were not normally distributed, accuracy was summarised using medians and compared between the four tools using Friedman tests followed by Dunn’s multiple-comparisons test. Accuracy–confidence differences were compared using a two-way repeated-measures ANOVA, with prediction tool as the repeated factor and protein category as the between-protein factor, followed by Tukey’s multiple-comparisons test between tools within each categoryand Šidák’s multiple-comparisons test between categories within each tool. Multiplicity-adjusted P-values below 0.05 were considered significant.

Fusion parent-region accuracy was compared between tools using a Friedman test followed by Dunn’s multiple-comparisons test, with each parent region treated as a matched set. Inter-tool similarity was compared between non-fusion and fusion proteins using a two-tailed Mann–Whitney test.

### Structure visualisation

Protein structures were visualised using ChimeraX (version 1.12 2026-06-12). Structures are shown at the same scale and and angle, after alignment using ChimeraX’s built-in matchmaker function. Residues of computationally-derived protein structures are colour-coded by ***B***-factor using the AlphaFold colour scale via color `bfactor all palette alphafold`.

## Results

### Performance benchmarking

#### Resource usage and runtime

Runtime and resource usage for the three tools across the 40 non-fusion proteins are summarised in Table 1. All three tools completed predictions for every protein. ESMFold was the fastest, with a median wall-clock time of 60.08 seconds per protein and 3.73 hours in total, compared with 17.45 hours for Boltz-2 and 62.23 hours for OmegaFold. ESMFold also used the least CPU time overall (103.80 CPU-hours, against 265.34 for Boltz-2 and 625.23 for OmegaFold). Median RAM use was similar across the three tools, ranging from about 7.3 to 9.3 GB per protein.

**Table 1.**
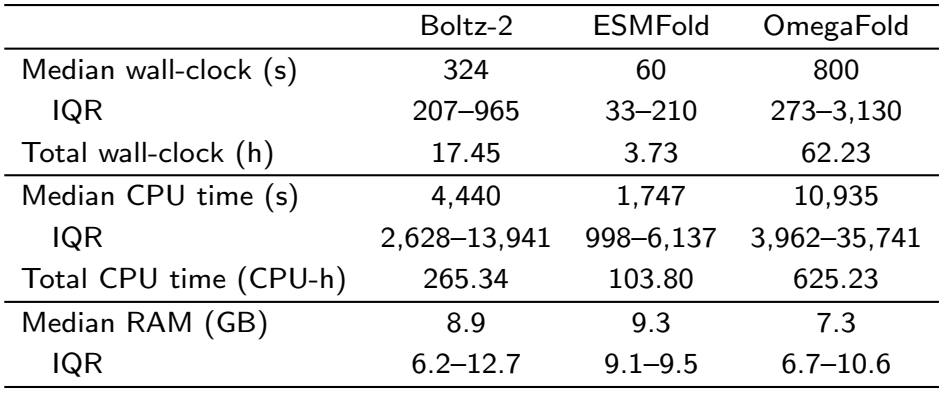
Runtime and resource usage for Boltz-2, ESMFold and OmegaFold across the 40 non-fusion proteins. Values are medians per protein, with interquartile range (IQR) shown underneath, and totals across all 40 proteins.

|  | Boltz-2 | ESMFold | OmegaFold |
|---|---|---|---|
| Median wall-clock (s) | 324 | 60 | 800 |
| IQR | 207–965 | 33–210 | 273–3,130 |
| Total wall-clock (h) | 17.45 | 3.73 | 62.23 |
| Median CPU time (s) | 4,440 | 1,747 | 10,935 |
| IQR | 2,628–13,941 | 998–6,137 | 3,962–35,741 |
| Total CPU time (CPU-h) | 265.34 | 103.80 | 625.23 |
| Median RAM (GB) | 8.9 | 9.3 | 7.3 |
| IQR | 6.2–12.7 | 9.1–9.5 | 6.7–10.6 |

Among the characterised control proteins, AlphaFold3 produced the highest median reference-normalised TM-score (0.986), followed by ESMFold (0.973), OmegaFold (0.966) and Boltz-2 (0.955). Pairwise differences were then assessed using Dunn’s multiple-comparisons test, with P-values adjusted for multiple testing. ESMFold produced significantly higher TM-scores than Boltz-2 (*P* = 0.0004). AlphaFold3 produced significantly higher TM-scores than OmegaFold (*P* = 0.0029) and Boltz-2 (*P <* 0.0001). No significant differences were detected for the remaining pairwise comparisons.

Among the cancer-associated proteins, AlphaFold3 produced the highest median reference-normalised TM-score (0.960), followed by OmegaFold (0.944), ESMFold (0.931) and Boltz-2 (0.918). Pairwise differences were then assessed using Dunn’s multiple-comparisons test, with P-values adjusted for multiple testing. AlphaFold3 produced significantly higher TM-scores than OmegaFold (*P* = 0.0132) and Boltz-2 (*P* = 0.0087). No significant differences were detected for the remaining pairwise comparisons.

Across the combined protein dataset, AlphaFold3 produced the highest median reference-normalised TM-score (0.977), followed by ESMFold (0.954), OmegaFold (0.952) and Boltz-2 (0.929). Pairwise differences were assessed using Dunn’s multiple-comparisons test, with P-values adjusted for multiple testing. ESMFold produced significantly higher TM-scores than OmegaFold (*P* = 0.0268) and Boltz-2 (*P* = 0.0006). AlphaFold3 produced significantly higher TM-scores than OmegaFold and Boltz-2 (*P <* 0.0001 for both comparisons). No significant differences were detected for the remaining pairwise comparisons.

Most characterised proteins received consistently high TM-scores across all four prediction tools. Carbonic anhydrase 2, haemoglobin alpha, superoxide dismutase 1, cytochrome C and lysozyme C were among the most accurately predicted proteins. Calmodulin-1 had the lowest mean TM-scores and the greatest variation between the four tools, ranging from 0.381 (Boltz-2) to 0.871 (ESMFold).This spread was reflected in which calcium-binding conformation/structure each tool had predicted (Table 2; Figure 6). Growth hormone showed substantial variation between tools, with AlphaFold3 producing a considerably higher TM-score than the other three models.

**Table 2.** Calmodulin-1 TM-scores against calcium-free and calcium-bound reference structures. Reference-normalised TM-scores for each tool against 1CFD (calcium-free) and 1CLL (calcium-bound).

| Tool | 1CLL (calcium-bound) | 1CFD (calcium-free) | Closer match |
|---|---|---|---|
| AlphaFold3 | 0.787 | 0.436 | Calcium-bound |
| Boltz-2 | 0.381 | 0.557 | Calcium-free |
| ESMFold | 0.871 | 0.436 | Calcium-bound |
| OmegaFold | 0.533 | 0.445 | Calcium-bound |

**Figure 1.**
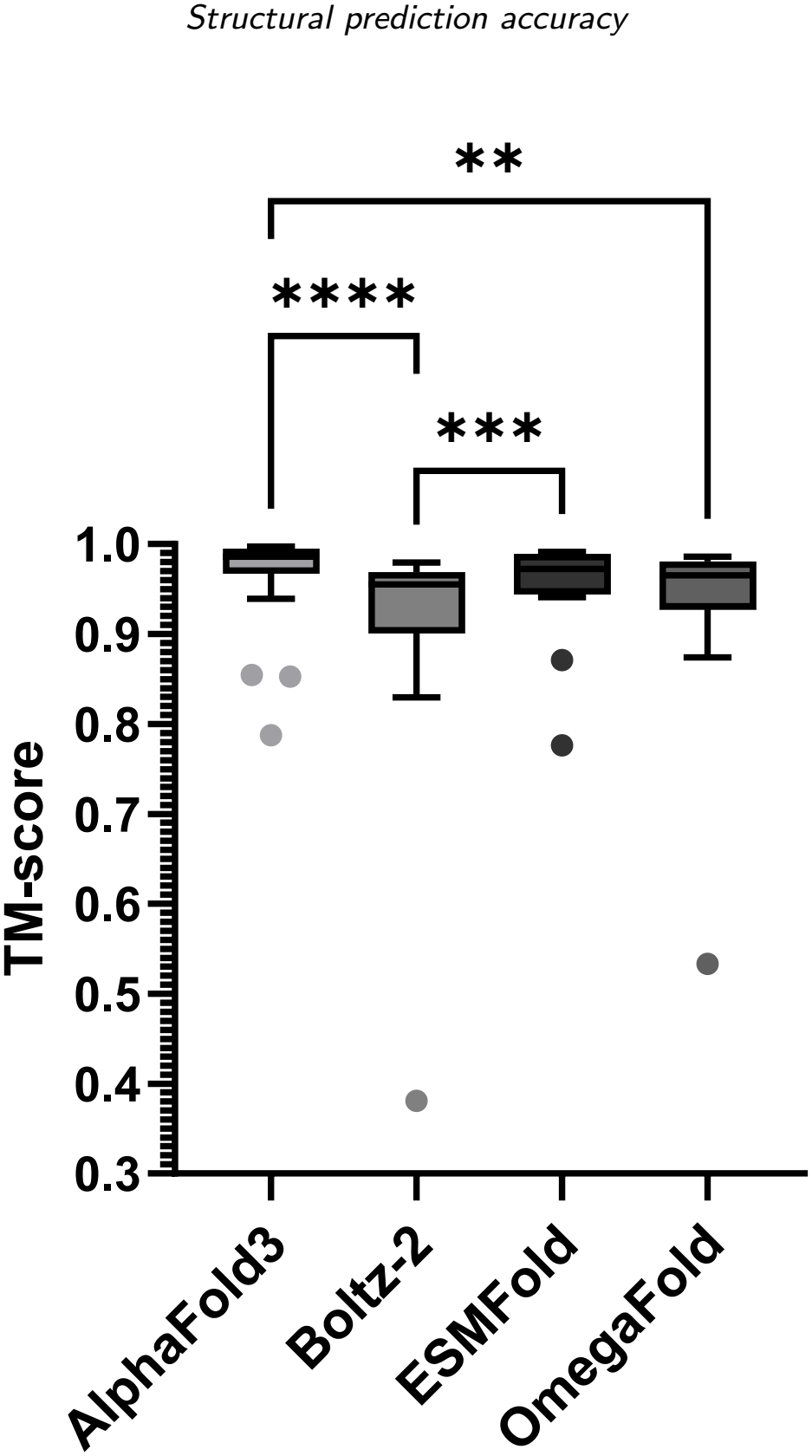
Structural prediction accuracy for characterised control proteins. Reference normalised TM-scores were compared across ESMFold, OmegaFold, Boltz-2 and AlphaFold3 for 18 proteins with validated experimental reference structures. Differences between tools were analysed using a Friedman test followed by Dunn’s multiple-comparisons test. Adjusted P-values are shown as (^*\*\**^*P <* 0.01), (^*\*\*\**^*P <* 0.001) and (^*\*\*\*\**^*P <* 0.0001).

**Figure 2.**
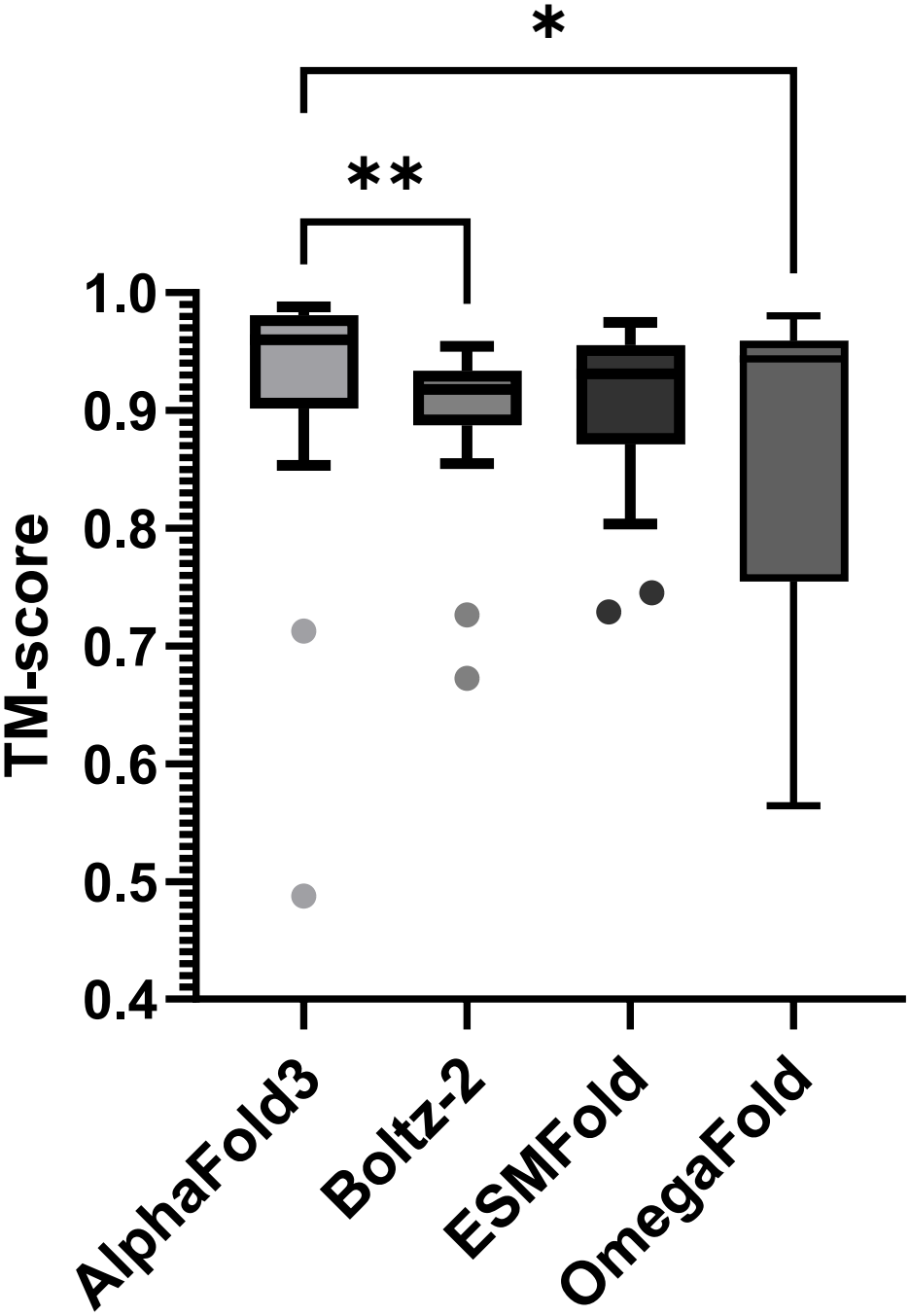
Structural prediction accuracy for cancer-associated proteins. Reference-normalised TM-scores were compared across ESMFold, OmegaFold, Boltz-2 and AlphaFold3 for 20 cancer-associated proteins with validated experimental reference structures. Differences between tools were analysed using a Friedman test followed by Dunn’s multiple-comparisons test. Adjusted P-values are shown as (^*\**^*P <* 0.05) and (^*\*\**^*P <* 0.01).

**Figure 3.**
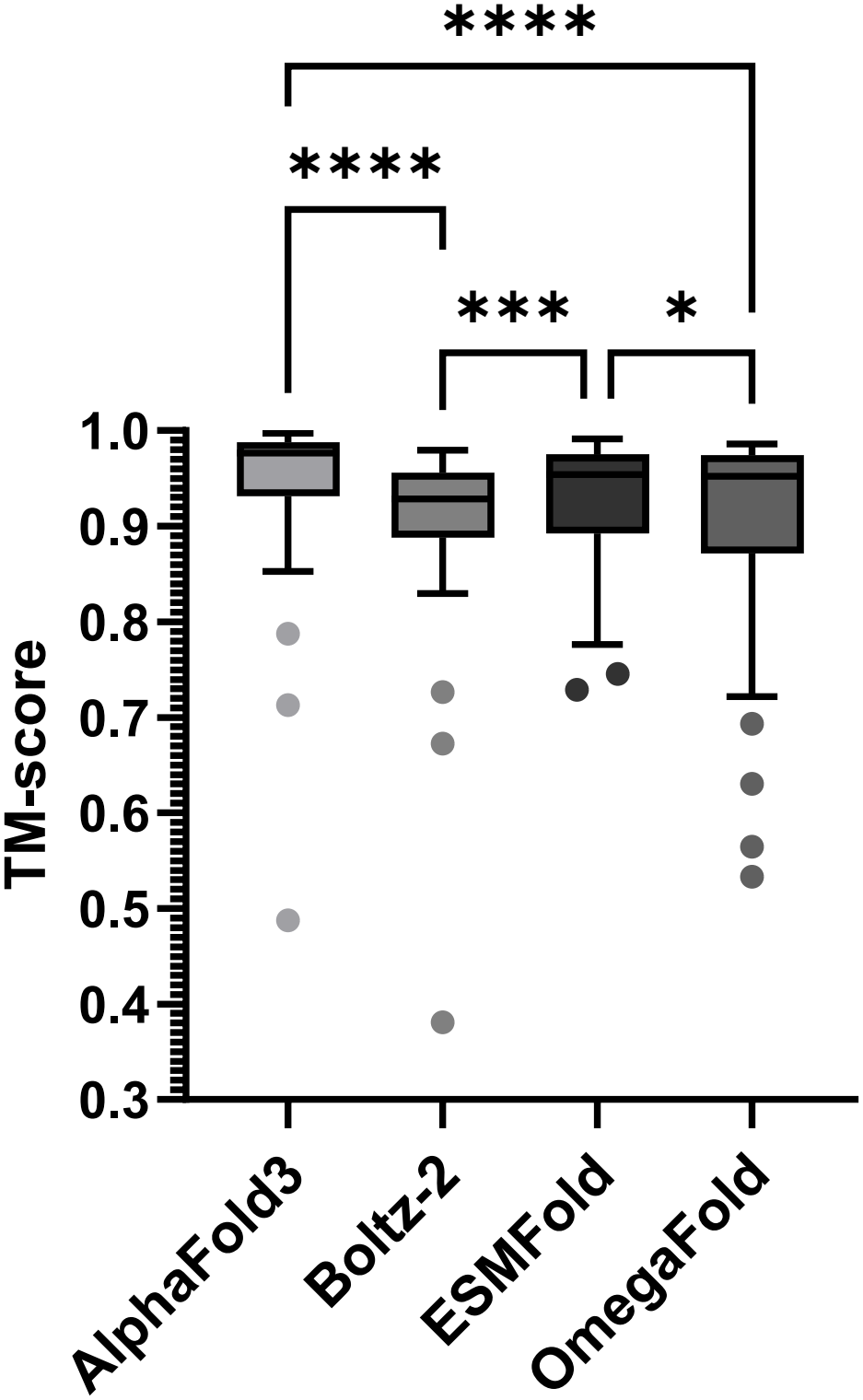
Overall structural prediction accuracy across characterised control and cancer-associated proteins. Reference-normalised TM-scores were compared across ESMFold, OmegaFold, Boltz-2 and AlphaFold3 for the combined set of 38 proteins with validated experimental reference structures. Differences between tools were analysed using a Friedman test followed by Dunn’s multiple-comparisons test. Adjusted P-values are shown as (^*\**^*P <* 0.05), (^*\*\*\**^*P <* 0.001) and (^*\*\*\*\**^*P <* 0.0001).

**Figure 4.**
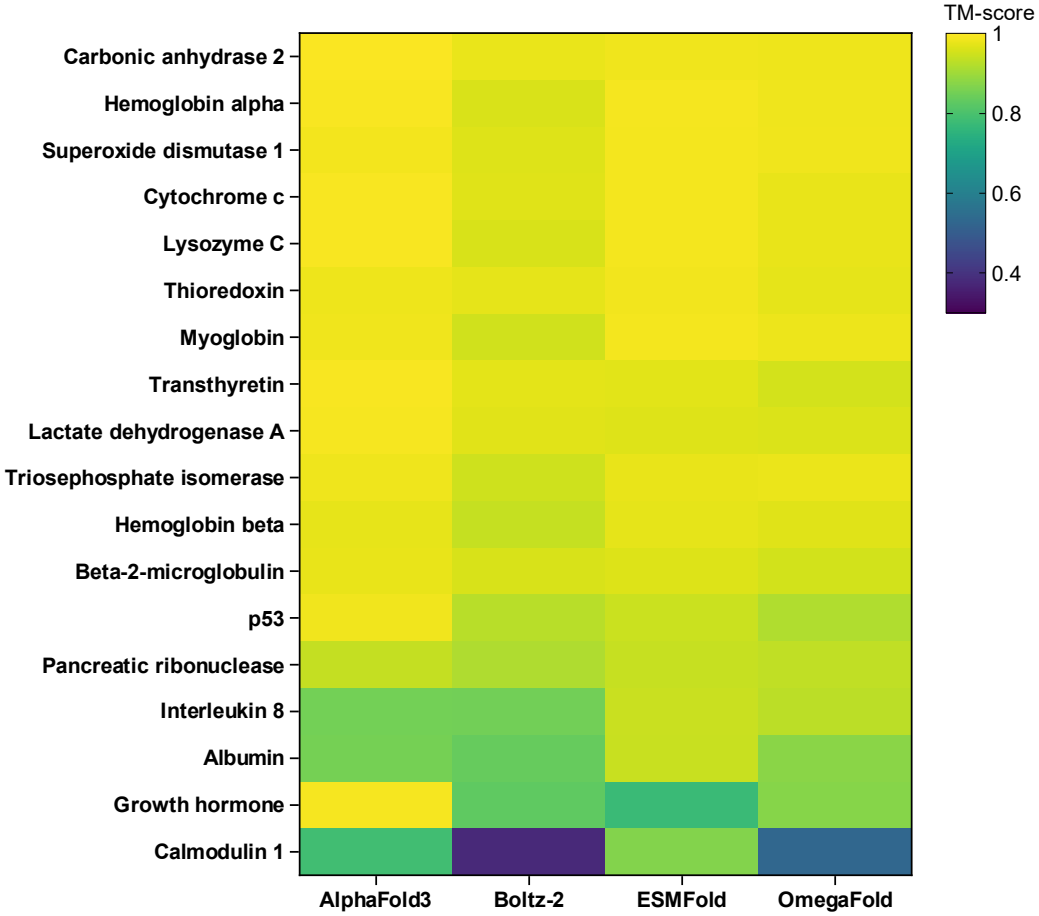
Structural prediction accuracy across characterised control proteins. Reference-normalised TM-scores are shown for 18 proteins predicted using ESMFold, OmegaFold, Boltz-2 and AlphaFold3. Proteins are ordered from highest to lowest mean TM-score across the four tools, with yellow indicating greater TM-score and purple indicating lower TM-score.

**Figure 5.**
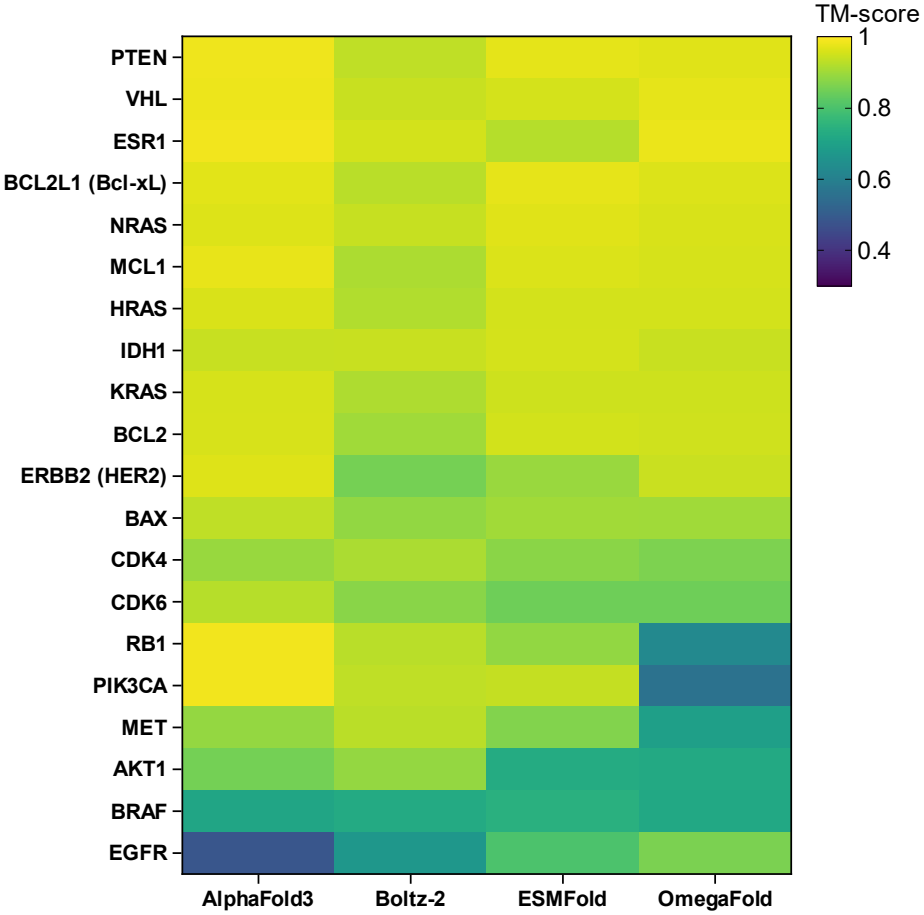
Structural prediction accuracy across cancer-associated proteins. Reference-normalised TM-scores are shown for 20 cancer-associated proteins predicted using ESMFold, OmegaFold, Boltz-2 and AlphaFold3. Proteins are ordered from highest to lowest mean TM-score across the four tools, with yellow indicating greater structural agreement with the experimental reference and purple indicating lower agreement.

**Figure 6.**
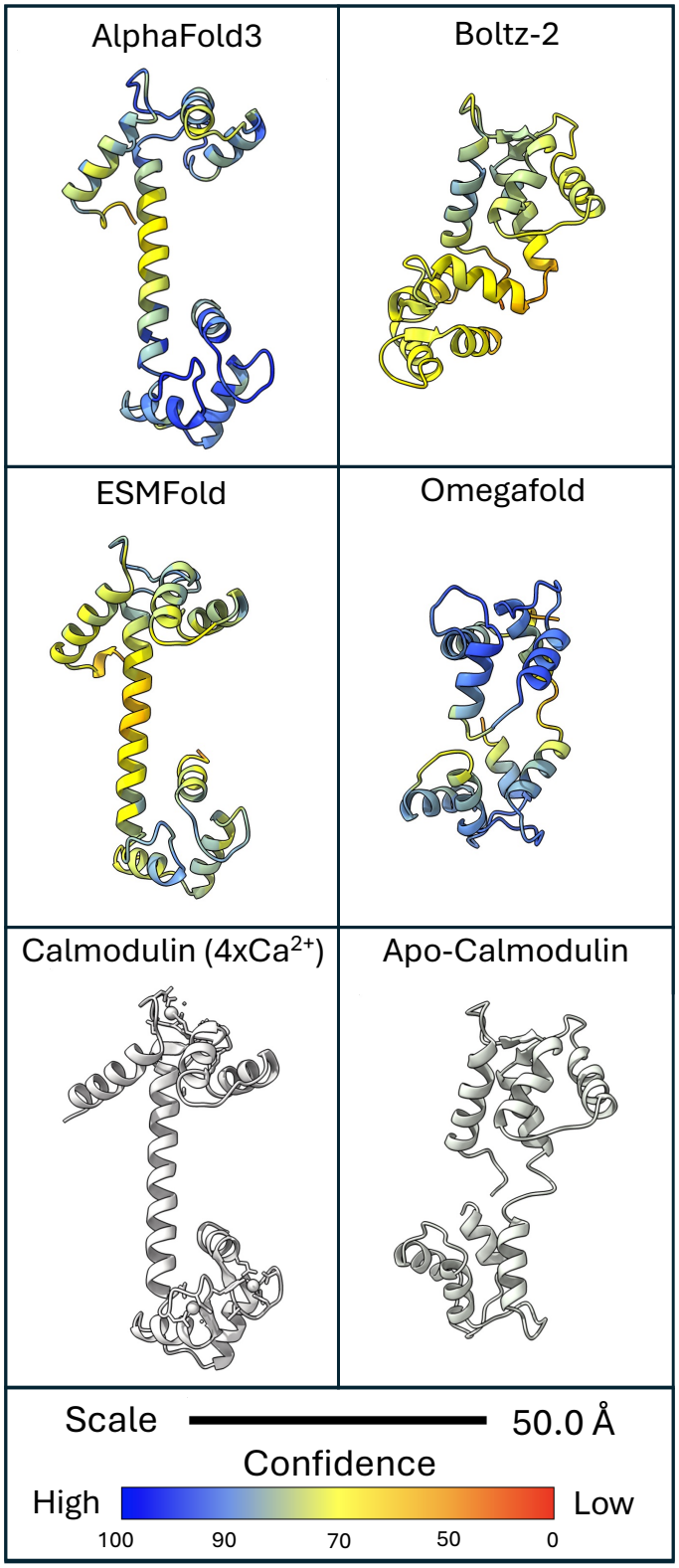
Predicted structure of human Calmodulin-1. Predictions from four tools, visualised in ChimeraX. Computationally-folded structures are coloured by confidence value, as shown on the key provided. Experimentally-determined structures of Calmodulin variants are shown in white. A 50 Angstrom (Å) scale is also provided.

**Figure 7.**
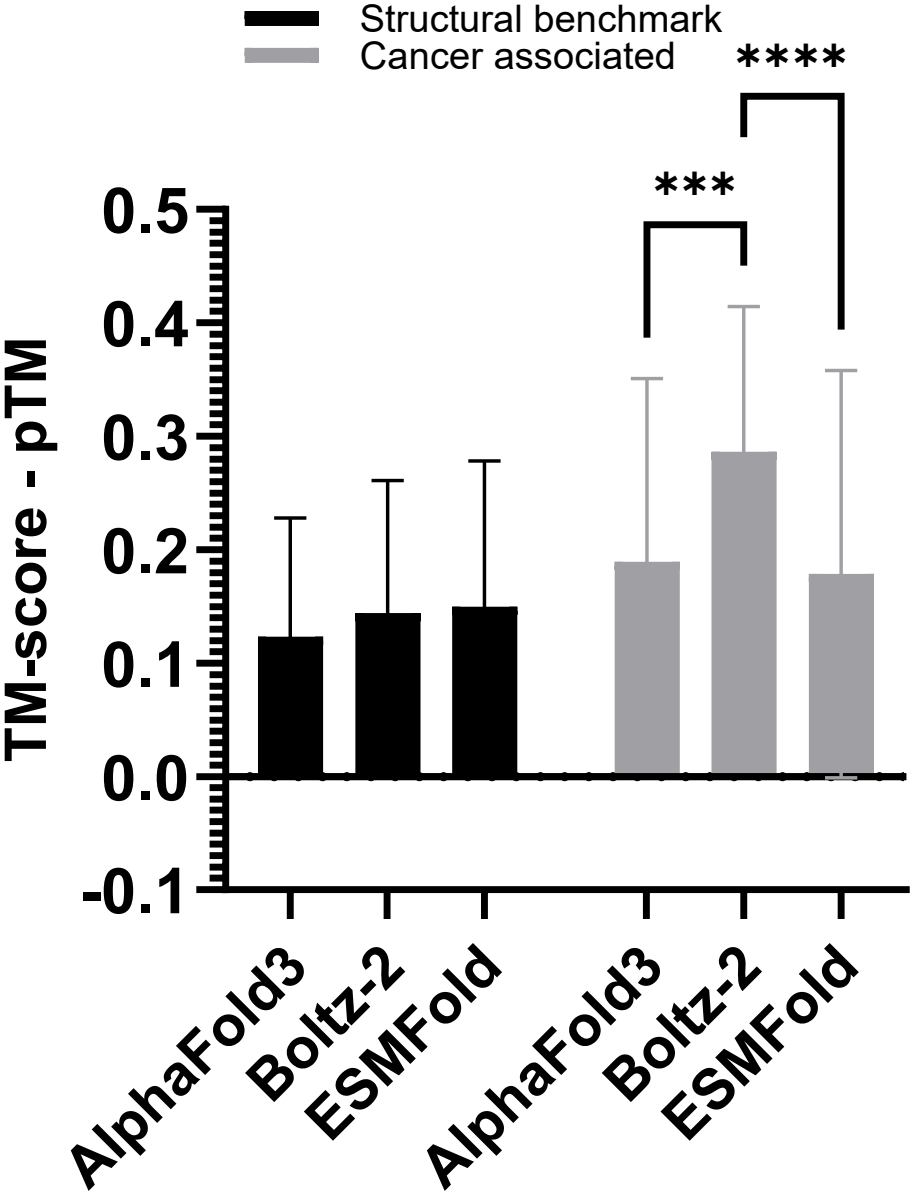
Accuracy–confidence differences across prediction tools and protein categories. The difference between the reference-normalised TM-score and model-reported pTM is shown for ESMFold, Boltz-2 and AlphaFold3 across characterised control proteins (n = 18) and cancer-associated proteins (n = 20). Positive values indicate that pTM underestimated measured structural accuracy. Bars show the mean *±* SD. Differences were analysed using a two-way repeated-measures ANOVA followed by Tukey’s multiple-comparisons test. Adjusted P-values are shown as (^*\*\*\**^*P <* 0.001) and (^*\*\*\*\**^*P <* 0.0001).

**Figure 8.**
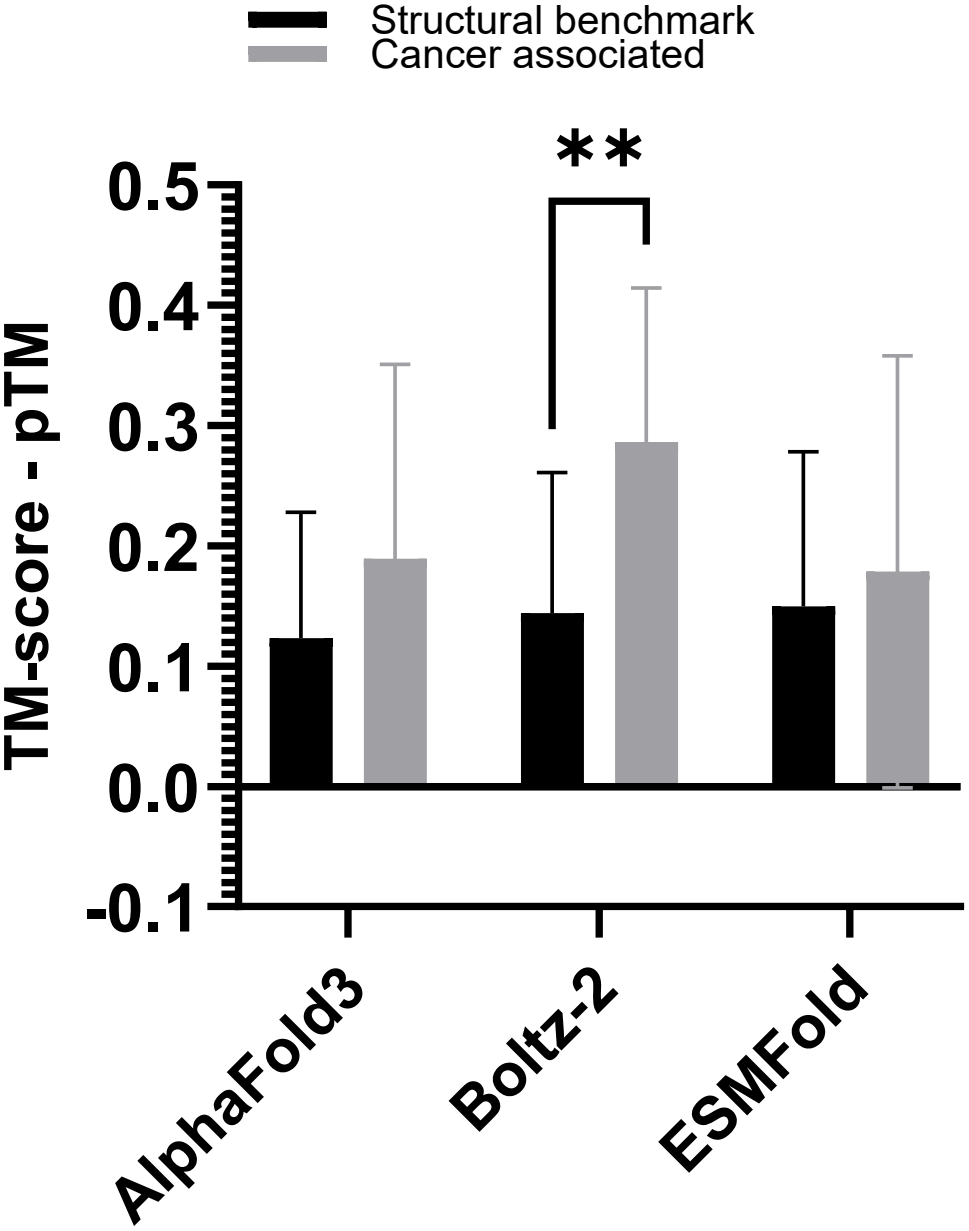
Comparison of accuracy–confidence differences between protein categories. The difference between the reference-normalised TM-score and model-reported pTM is shown for characterised control proteins (n = 18) and cancer-associated proteins (n = 20). Positive values indicate that pTM underestimated measured structural accuracy. Bars show the mean *±* SD. Differences were analysed using a two-way repeated-measures ANOVA followed by Šídák’s multiple-comparisons test. The adjusted P-value is shown as (^*\*\**^*P <* 0.01).

PTEN, VHL, ESR1 and BCL2L1 received consistently high TM-scores across the four prediction tools. EGFR had the lowest mean TM-score, followed by BRAF and AKT1. Greater variation between tools was observed for RB1, PIK3CA and EGFR. OmegaFold produced notably lower scores for RB1 and PIK3CA than the other tools, whilst AlphaFold3 produced the lowest score for EGFR. CDK4 was scored against an X-ray structure of CDK4 bound to cyclin D3 (3G33), against which all four tools achieved TM-scores between 0.864 and 0.913. The originally retrieved reference, an 8 Å cryo-EM structure of CDK4 bound to the HSP90–CDC37 chaperone, did not meet the reference criteria; results against it are given in Supplementary Table S2.

#### Effect of reference conformation on calmodulin-1

Calmodulin-1 undergoes a large conformational change upon calcium binding (Chou et al., 2001). The apparent accuracy of each prediction depended strongly on which experimental conformation was used as the reference (Table 2). ESMFold, AlphaFold3 and OmegaFold more closely resembled the calcium-bound structure (1CLL), with TM-scores against 1CLL exceeding those against calcium-free calmodulin (1CFD) by 0.435, 0.351 and 0.088 respectively. In contrast, Boltz-2 more closely resembled the calcium-free conformation (TM-score 0.557 against 1CFD versus 0.381 against 1CLL). None of the tools was provided with calcium ions, so these differences reflect the conformation each model reproduced from the given, sequence alone.

#### Self-assessed confidence accuracy

Mean accuracy-confidence differences were almost all positive for the three models in both protein categories, indicating an overall tendency to underestimate structural accuracy. Among the characterised control proteins, no significant differences between models were detected (AlphaFold3, 0.123; Boltz-2, 0.144; ESMFold, 0.150). Among the cancer-associated proteins, Boltz-2 showed a significantly greater difference (0.287) than both ESMFold (0.186, adjusted *P <* 0.0001) and AlphaFold3 (0.189, adjusted *P* = 0.0004). No other pairwise differences between models were significant. The analysis identified a significant main effect of prediction tool (*P* = 0.0004) and a significant interaction between prediction tool and protein category (*P* = 0.0026).

The mean accuracy-confidence difference was greater for the cancer-associated proteins than for the characterised control proteins across all three tools. This difference was significant for Boltz-2, increasing from 0.144 in the control group to 0.287 in the cancer-associated group (adjusted *P* = 0.0029). No significant difference between protein categories was detected for ESMFold or AlphaFold3. The overall main effect of protein category was not significant (*P* = 0.065), although a significant interaction between prediction tool and protein category was detected (*P* = 0.0026).

#### Visual structure assessment

The structures of the two poorest-performing cancer-associated proteins, BRAF and EGFR, showed drastically different results for each of the four tools. In the case of BRAF (Figure 9), AlphaFold3, ESMFold, and Omegafold all produced substantially less compact structures than the reference protein, which contained a mixture of high-confidence regions of some consistency and disordered (confidence 50%) regions of no overlap. Boltz-2 instead produced an excessively compact protein with no clear high-confidence regions. For EGFR (Figure 10), this pattern largely held, with tools producing a mixture of similar high-confidence regions and disordered segments. Consistent regions are similar in structure to the beta-sheet regions present in the EGFR reference structure; however, in this case, all tools also produced a long linear alpha-helix region that is wholly absent from the reference.

**Figure 9.**
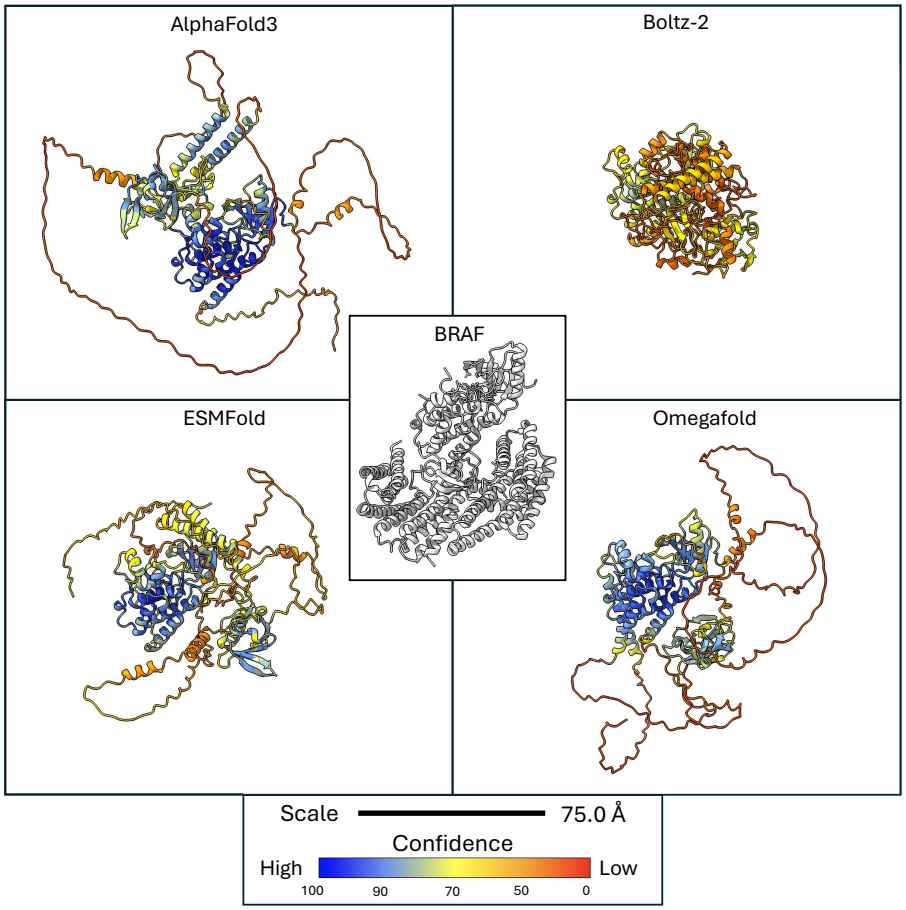
Predicted structure of the cancer-associated protein BRAF. Predictions from four tools, visualised in ChimeraX. Computationally-folded structures are coloured by confidence value, as shown on the key provided. Experimentally-derived reference structure is shown in white. A 75 Angstrom (Å) scale is also provided.

**Figure 10.**
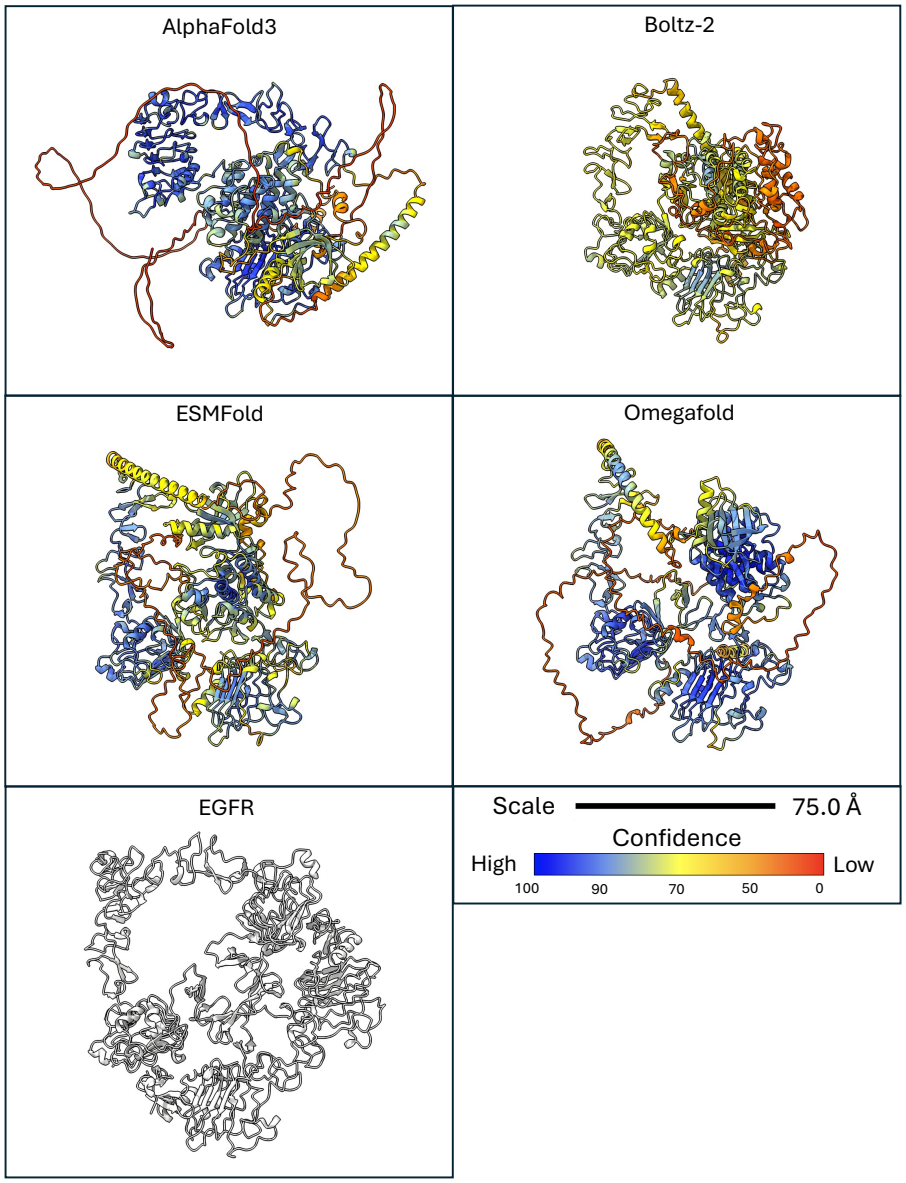
Predicted structure of the cancer-associated protein EGFR. Predictions from four tools, visualised in ChimeraX. Computationally-folded structures are coloured by confidence value, as shown on the key provided. Experimentally-derived reference structure is shown in white. A 75 Angstrom (Å) scale is also provided.

### Performance on novel chimeric structures vs. known proteins

#### Accuracy against parent reference structures

No experimental structures exist for the fusion proteins, so prediction accuracy was assessed against the experimental structures of the parent proteins. Only parents sharing at least 90% of their sequence with the fusion were used, giving 12 parent regions from 11 fusion proteins (6 Parent A, 6 Parent B).

Among the fusion parent regions, AlphaFold3 produced the highest median reference-normalised TM-score (0.844), followed by OmegaFold (0.828), ESMFold (0.802) and Boltz-2 (0.714). Accuracy differed significantly between tools (Friedman test, *P* = 0.0017). Pairwise differences were then assessed using Dunn’s multiple-comparisons test, with P-values adjusted for multiple testing. AlphaFold3 (*P* = 0.0094) and OmegaFold (*P* = 0.0160) produced significantly higher TM-scores than Boltz-2. No significant differences were detected for the remaining pairwise comparisons, and AlphaFold3 and OmegaFold additionally, performed almost identically (*P >* 0.9999).

Scores fell into two clear groups rather than a continuous spread (Figure 11). Most parent regions were predicted with high accuracy by every tool, while BCR-ABL1 (A), CBFB-MYH11 (A), PML-RARA (B), SS18-SSX1 (A) and SS18-SSX2 (A) scored below 0.52 with all four tools. Parent B regions generally scored higher than Parent A regions (median of tool means 0.909 vs 0.487), although there were exceptions in both directions, including RUNX1-RUNX1T1 (A) at 0.917 and PML-RARA (B) at 0.350.

**Figure 11.**
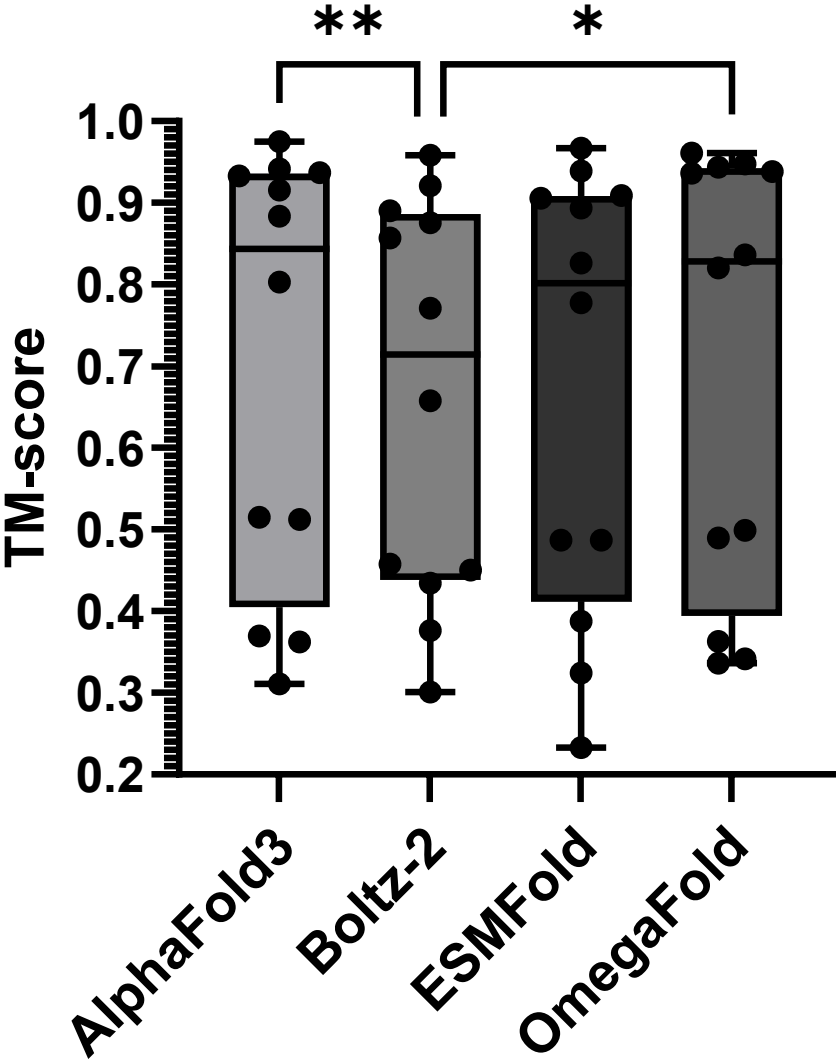
Structural prediction accuracy for fusion protein parent regions. Reference-normalised TM-scores were compared across AlphaFold3, Boltz-2, ESMFold and OmegaFold for 12 parent regions from 11 fusion proteins, using parent structures sharing at least 90% of their sequence with the fusion. Boxes show the median and interquartile range, whiskers show the minimum to maximum and points show individual parent region scores. Differences between tools were analysed using a Friedman test followed by Dunn’s multiple-comparisons test. Adjusted P-values are shown as (^*\**^*P <* 0.05) and (^*\*\**^*P <* 0.01).

**Figure 12.**
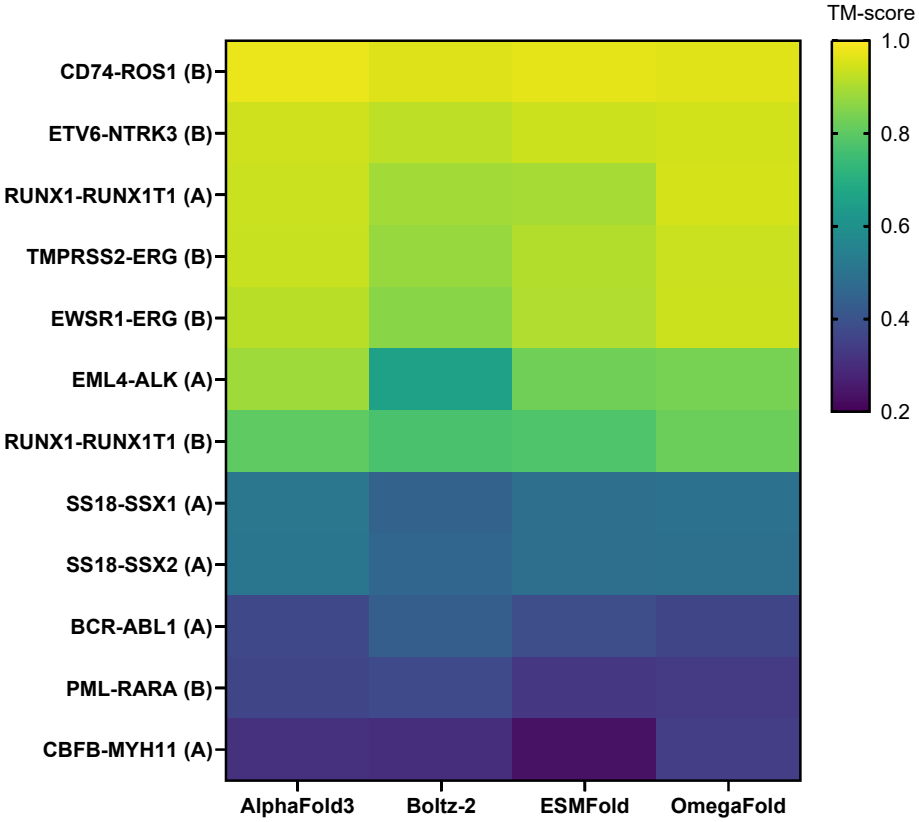
Structural prediction accuracy across fusion protein parent regions. Reference-normalised TM-scores are shown for 12 parent regions from 11 fusion proteins predicted using AlphaFold3, Boltz-2, ESMFold and OmegaFold. (A) and (B) indicate whether the region corresponds to Parent A or Parent B. Regions are ordered from highest to lowest mean TM-score across the four tools, with yellow indicating greater TM-score and purple indicating lower TM-score.

#### Pairwise comparison of structural prediction

To assess whether the four tools converged on the same structure, each pair of predicted models for every protein was compared using US-align, and the pairwise TM-scores were averaged to give a single inter-tool similarity score per protein. Inter-tool similarity was substantially lower for the fusion proteins (median 0.272, n = 20) than for the characterised and cancer-associated proteins (median 0.853, n = 40; Mann-Whitney U = 81, *P <* 0.0001) (Figure 13). Only DNAJB1-PRKACA reached a level of agreement typical of the characterised proteins (0.898), while 11 of the 20 fusions scored below 0.30.

**Figure 13.**
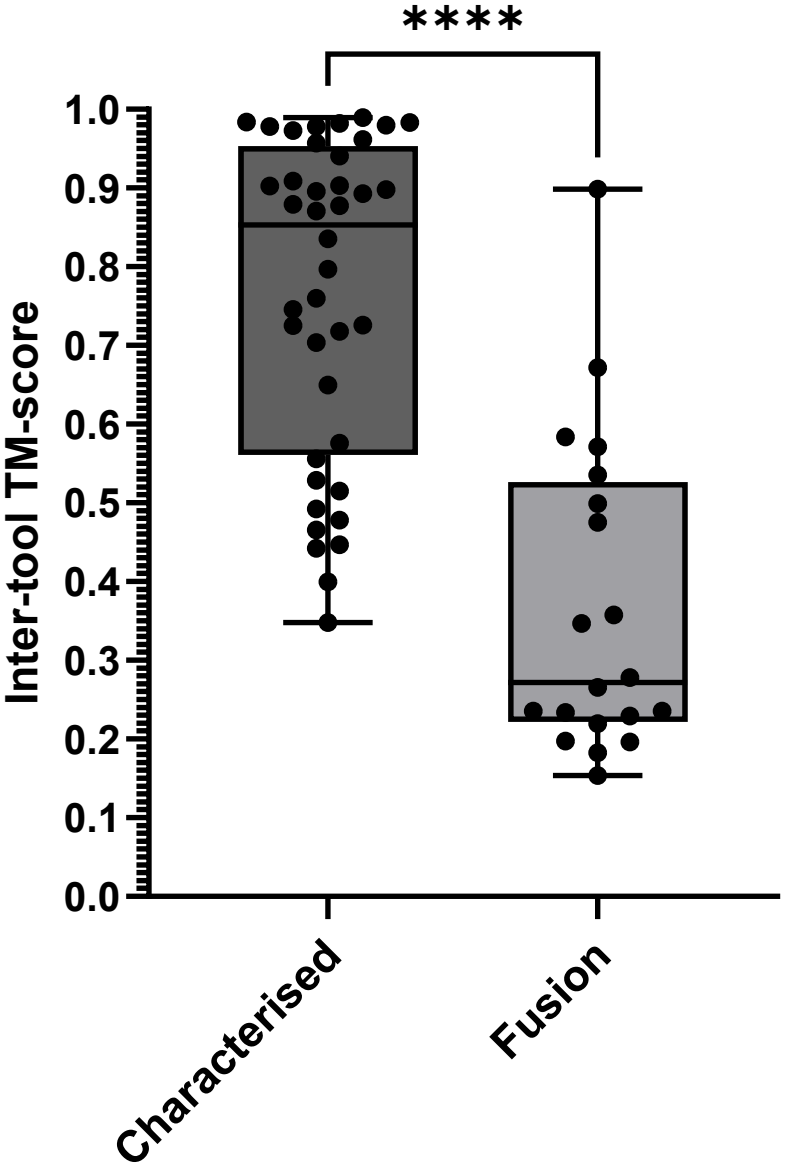
Agreement between prediction tools for characterised and fusion proteins. For each protein, full-length predicted structures from AlphaFold3, Boltz-2, ESMFold and OmegaFold were compared pairwise using US-align, and the resulting TM-scores were averaged to give one inter-tool similarity score per protein. Scores are shown for 40 characterised and cancer-associated proteins and 20 fusion proteins. Boxes show the median and interquartile range, whiskers show the minimum to maximum and points show individual proteins. The groups were compared using a two-tailed Mann-Whitney test (^*\*\*\*\**^*P <* 0.0001).

#### TMPRSS2-ERG

In the case of TMPRSS2-ERG, all four tools produced vastly different structures, with Boltz-2 notably producing a far more dense and compact structure versus the other tools, with lower confidence scores assigned (Figure 14). Most residues overall ranged from low confidence to disorder. Despite this, all four tools produced a functional domain capturing the structure of the parent protein ERG, which is the primary driver of oncogenic activity (Sekar et al., 2026). Beyond this, the tools other than Boltz-2 returned other consistent regions, notably a second high-confidence domain and an extended straight alpha-helix region.

**Figure 14.**
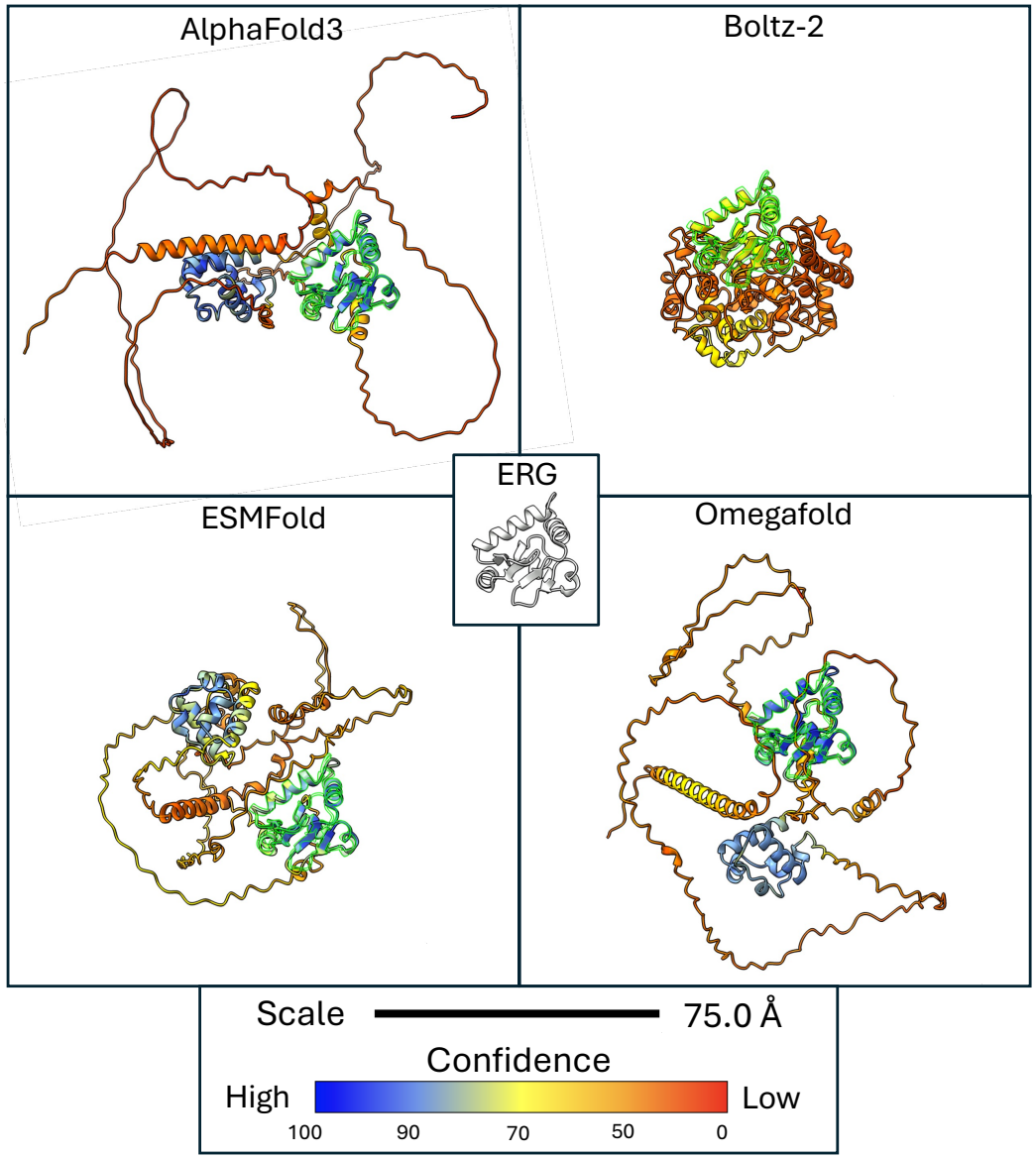
Predicted structure of the cancer fusion protein TMPRSS2-ERG. Predictions from four tools, visualised in ChimeraX. Computationally-folded structures are coloured by confidence value, as shown on the key provided.. Green outlines show the best alignment of the active-site parent protein ERG, with a reference structure shown in white. A 75 Angstrom (Å) scale is also provided.

#### DNAJB1-PRKACA

Given the raw amino acid sequence of DNAJB1-PRKACA, all four tools produced highly similar results, with self-assessed confidence for 90% for the majority of residues; however, the parent protein PRKACA, which contributes the fusion protein’s functional kinase domain (Bolger, 2022), aligned poorly and inconsistently to all resulting structures (Figure 15). Though the sequence of PRKACA is almost fully contained within the fusion protein, this did not extend to the structure of PRKACA. The expected beta-sheets that form its experimentally-determined structure are wholly absent from all tools’ results.

**Figure 15.**
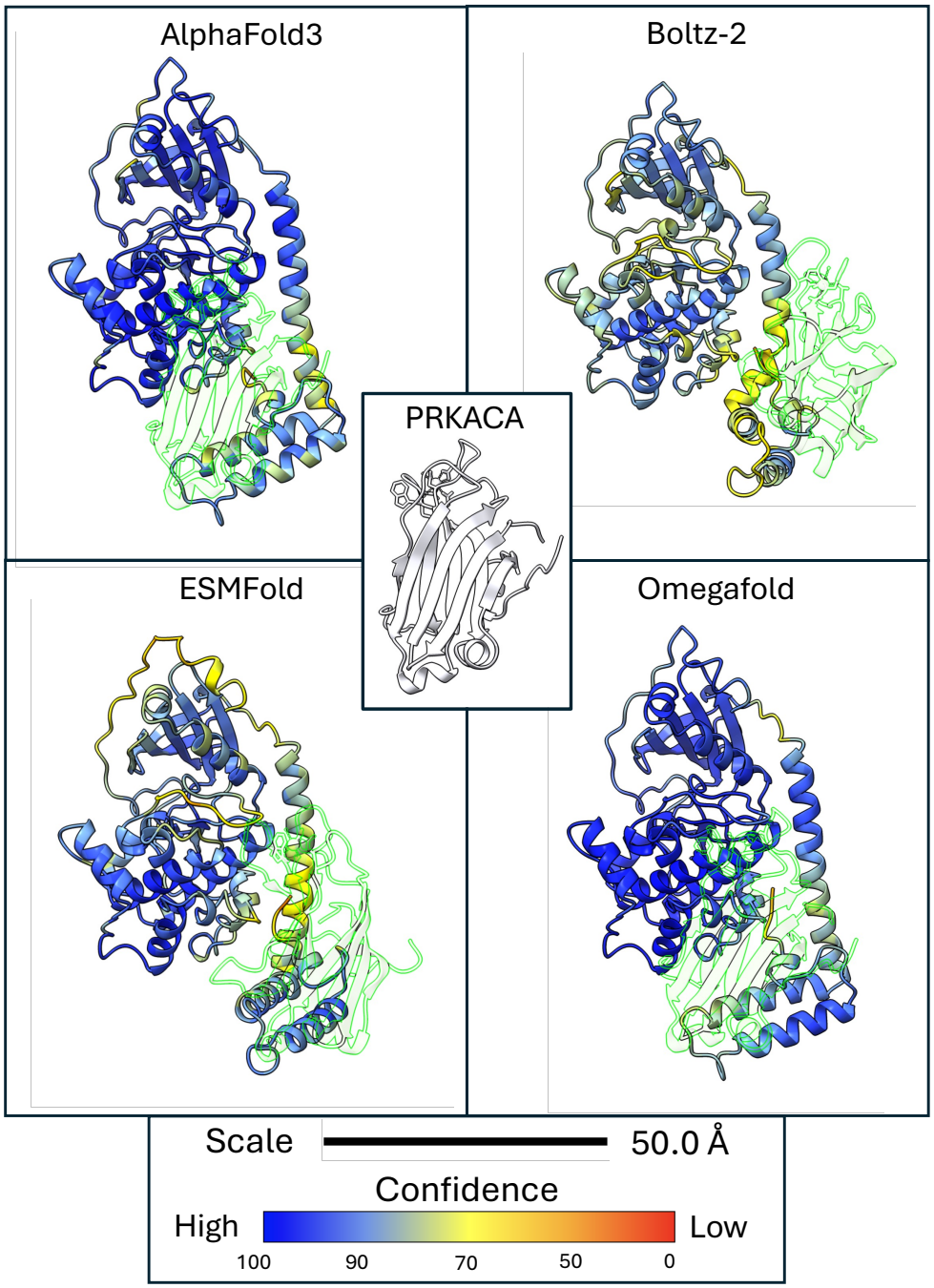
Predicted structure of the cancer fusion protein DNAJB1-PRKACA. Predictions from four tools, visualised in ChimeraX. Computationally-folded structures are coloured by confidence value, as shown on the key provided. Green outlines show the best alignment of the active-site parent protein PRKACA, with a reference structure shown in white. A 50 Angstrom (Å) scale is also provided.

## Discussion

### Prediction reliability

#### Overall accuracy

All four tools predicted the characterised control proteins with high accuracy, with median TM-scores above 0.95, and AlphaFold3 had the highest median TM-score for both the characterised and cancer-associated proteins. However, the differences between tools were minor. Across the 38 proteins, median TM-scores ranged from 0.929 for Boltz-2 to 0.977 for AlphaFold3, so all four tools would usually recover, broadly, the correct overall fold of a well-characterised protein.

One caveat applies to all of these results. The tools were trained on structures from the Protein Data Bank, and many of the reference structures used here, or close homologues of them, are likely to have been in their training data. The high accuracy on characterised proteins may therefore partly reflect structures the models have already seen, and probably overestimates how they would perform on genuinely new proteins. This is one reason the fusion proteins comparison is crucial, as their full-length structures have never been solved.

#### Calmodulin

The case of Calmodulin-1 demonstrates a failure mode of computational protein folding, where a more common structure which is not the best fit for the input data may be returned in place of a rarer but technically more accurate structure. Three of the four folding methods tested (excluding Omegafold) allow explicit specification of atoms and small molecules to include for co-folding with the protein. A strictly accurate result in this case would be folds specifying the sequence of calmodulin only returning the apo-calmodulin configuration, while those additionally specifying four or more calcium ions returning the calmodulin (4xCa2+) configuration. This may not be desirable in all cases or to all users, however, as it potentially requires the input of much more detailed information to produce common structures that may be widely accepted.

#### Model confidence

For ESMFold, Boltz-2 and AlphaFold3, pTM underestimated measured accuracy in almost every case, so the tools were conservative rather than overconfident. A likely explanation is that pTM is calculated over the whole predicted chain, including flexible tails and linkers, whereas TM-score here was calculated only over the region resolved in the experimental structure. Taken together, a low pTM should not be taken to mean that the folded core of a prediction is wrong.

### Performance on cancer-related proteins

Accuracy was lower for the cancer-associated proteins than for the characterised control proteins, although AlphaFold3 still achieved a median TM-score of 0.960. The lowest-scoring proteins, including EGFR, BRAF and AKT1, are large multi-domain signalling proteins, and EGFR and BRAF also had among the lowest agreement between tools of any non-fusion protein (mean inter-tool TM-scores of 0.45 and 0.40). The gap between measured accuracy and pTM was also larger for the cancer-associated proteins, particularly for Boltz-2, where it doubled from 0.144 to 0.287. Notably, BRAF contains two separate functional regions connected by a flexible hinge and has both C- and N-terminal extensions (Ducreux et al., 2019). This fits the explanation above, as larger proteins with more flexible regions would lower pTM without affecting the accuracy of their folded domains. EGFR, on the other hand, has numerous conformational states depending on its state as a monomer, dimer, ligand binding, membrane binding, and others (Yoon et al., 2026). In this case, the issue may be more similar to that of Calmodulin-1, where a range of possible structures causes tools to return an averaged approximation. CDK4 showed how much the choice of reference can matter. Its scores rose to 0.864–0.913 once a chaperone-bound cryo-EM structure was replaced with an X-ray structure. As with calmodulin, a TM-score measures agreement with one particular experimental structure, and the choice of reference carries similar value to the choice of tool.

### Performance on novel and chimeric proteins

When predictions were compared with the parent proteins, accuracy split into two groups: most parent regions were predicted well by every tool, while five regions scored below 0.52 with all four. AlphaFold3 and OmegaFold performed best, and Boltz-2 was significantly worse than both, in line with its lower scores on the non-fusion proteins. Parent B regions generally scored higher than Parent A regions, which may reflect that the 3^*′*^ partner often contributes the folded functional domain, such as a kinase, although with six regions per group this could not be tested statistically.

The four tools also agreed with each other far less for the fusion proteins (median inter-tool TM-score 0.272) than for the characterised and cancer-associated proteins (0.853). Among the non-fusion proteins, the lowest agreement was seen for proteins with large disordered or flexible regions, including p53, BRAF, EGFR and polyubiquitin-C, which suggests that disagreement between tools is driven mainly by regions without a single stable fold. Fusion proteins appear to contain much more of this.

The fusions selected to capture the range of possible results, <u>TMPRSS2-ERG</u> and <u>DNAJB1-PRKACA,</u> show vastly different and potentially counterintuitive results. The first, TMPRSS2-ERG, which produces structures that appear inconsistent by alignment-based metrics, likely successfully capture the most biologically-important structures of the fusion protein, with the structure of the ERG transcription factor clearly and confidently returned by all methods. While Boltz-2 produced a much more dense structure, all others showed regions of consistency separated by long disordered regions. Thus, low alignment scores likely primarily resulted from the flexibility of poor-confidence or intrinsically-disordered regions, whereas functional domains were calculated at higher confidence and were largely consistent. In the case of DNAJB1-PRKACA, the opposite effect was found. All four structures produced were nearly identical, but none contained any domain resembling the parent protein PRKACA, which must be present for the fusion’s known mode of action. This demonstrates that structural alignment is likely a poor method of assessing the consistency of predicted proteins. It may be best to develop a method that instead aligns high-confidence domains that are most likely to hold functional roles, giving a higher weighting to these; however, this does not address the issue of proteins that do consistently produce high-confidence folds over their entirety, but do not produce the domains necessary for that expected function. In cases such as DNAJB1-PRKACA, it remains unclear if the software predictions are incorrect in a uniform way, or if the biology of this protein is different than what is expected.

More generally, each tool returns a single structure, and for regions that lack one stable fold, that structure seems to be shaped as much by the model’s own tendencies as by the sequence, as seen in Boltz-2’s compact predictions. Future tools would benefit from representing this uncertainty more explicitly, for example by predicting a range of conformations rather than a single model.

### Limitations

This study used a modest number of proteins, particularly for the fusion parent-region analysis, which included only 12 regions. The fusion results should therefore be treated as a first look rather than a definitive comparison. Accuracy was measured with TM-score only, which reflects the overall fold rather than local detail, and no ligands, ions or binding partners were given to the tools. ESMFold, OmegaFold and Boltz-2 were run on CPU and AlphaFold3 on GPU, so AlphaFold3’s resource use could not be compared directly with the other three tools. The study also assesses a small number of proteins overall; however, each was critically evaluated on an individual basis, allowing the observation of patterns that are washed out in larger purely-statistical comparisons. The study also lacks truth data for the fusion proteins tested. This is unfortunately out of necessity, as nearly all proteins for which experimental structural predictions are publicly available are included in the training set of the models tested.

## Conclusions

In terms of assessed accuracy alone, AlphaFold3 performed best of the four tools tested. In cases where GPU resources are not available, AlphaFold3 pre-trained models cannot be used, or faster run speed is a requirement, ESMFold was largely equivalent in accuracy to AlphaFold3. Boltz-2 and OmegaFold performed notably worse on a number of the sample proteins tested, with Boltz-2 observed to frequently produce unusually-dense folds in cases of low confidence. Additionally, OmegaFold lacked some of the features of other software, most notably in self-assessed accuracy and in the support for atom and small-molecule co-folding.

The findings of this research largely support that the protein-folding methods tested, while not perfectly reliable, are largely effective tools for determining structure from sequence alone. Rather that improvement to the tools themselves, however, we note that improvements are needed in methods of structural comparison to *assess* the results. Furthermore, subsequent generations of tools likely need to focus on differentiation of protein regions that can be conclusively folded, those which likely have deterministic fold-structure that the software is unable to determine, and those which are intrinsically disordered in nature. As newer approaches more effectively incorporate protein dynamics and structural flexibility, methods of protein visualisation and comparison that properly account for this information will be needed.

Finally, we note the importance of biological knowledge in assessing protein-folding results. The metrics used in this study, which included alignment scoring and self-assessed accuracy, largely fail to capture the biological plausibility of the structures returned. This presents a serious barrier to high-throughput assessment of novel protein structures such as in drug discovery. As shown for some of the protein cases evaluated in this paper, a human expert would be required to individually assess outputs and review the literature to properly interpret results.

### Future directions

In the future, we hope to test evaluate methodologies of both protein folding that explicitly account for conformational change and flexibility as well as alignment tools that weight these regions vs. more restricted functional domains appropriately. Should the cancer fusion proteins evaluated in this study be experimentally characterised, we would then revisit our assessment of these and determine if our original evaluations were accurate. Lastly, there may be opportunities for the development of AI-enabled workflows to support this assessment beyond what can be captured by quantitative metrics alone, allowing per-protein “critical review” to be performed at scale.

## Supplementary Information

### Data availability

Reference protein structures used are freely and publicly available from RCSB Protein Data Base: https://www.rcsb.org/. All scripts, sequences, predicted structures, reference manifests, and supplementary tables are available at https://github.com/CiaranMcGettrick/protein-structure-prediction-benchmarking.

### Supplementary tables

Supplementary Tables S1–S5 are provided in the accompanying file `Supplementary_Tables.xlsx`. Table S1 lists the experimental reference structures used for the non-fusion proteins; Table S2 gives TM-scores for all four tools against the originally retrieved references; Table S3 gives the final per-protein TM-scores for each tool; Table S4 shows parent associations and parent-region alignment scores; Table S5 shows inter-tool agreement for the 60 proteins tested.

## Acknowledgements

Research on this project was partially funded by a University of Galway School of Mathematical & Statistical Sciences Summer Studentship award. Analyses performed utilised CPU and GPU resources of the School of Mathematical & Statistical Sciences’s ***LUGH*** HPC system.

## Declarations

Some scripts were written with assistance from Claude Opus 5.5 (Anthropic). All code was reviewed and tested by the authors, who designed the study, curated the datasets, performed the analyses and interpreted the results.

The authors declare no conflicts of interest or other ethical issues related to this research.

## References

Josh Abramson, Jonas Adler, Jack Dunger, Richard Evans, Tim Green, Alexander Pritzel, Olaf Ronneberger, Lindsay Willmore, Andrew J. Ballard, Joshua Bambrick, Sebastian W. Bodenstein, David A. Evans, Chia-Chun Hung, Michael O’Neill, David Reiman, Kathryn Tunyasuvunakool, Zachary Wu, Akvilė Žemgulytė, Eirini Arvaniti, Charles Beattie, Ottavia Bertolli, Alex Bridgland, Alexey Cherepanov, Miles Congreve, Alexander I. Cowen-Rivers, Andrew Cowie, Michael Figurnov, Fabian B. Fuchs, Hannah Gladman, Rishub Jain, Yousuf A. Khan, Caroline M. R. Low, Kuba Perlin, Anna Potapenko, Pascal Savy, Sukhdeep Singh, Adrian Stecula, Ashok Thillaisundaram, Catherine Tong, Sergei Yakneen, Ellen D. Zhong, Michal Zielinski, Augustin Žídek, Victor Bapst, Pushmeet Kohli, Max Jaderberg, Demis Hassabis, and John M. Jumper. Accurate structure prediction of biomolecular interactions with alphafold 3. Nature, 630(8016):493–500, 2024. ISSN 1476-4687. doi: 10.1038/s41586-024-07487-w. URL https://doi.org/10.1038/s41586-024-07487-w.

David R Armstrong, John M Berrisford, Matthew J Conroy, Aleksandras Gutmanas, Stephen Anyango, Preeti Choudhary, Alice R Clark, Jose M Dana, Mandar Deshpande, Roisin Dunlop, Paul Gane, Romana Gáborová, Deepti Gupta, Pauline Haslam, Jaroslav Koča, Lora Mak, Saqib Mir, Abhik Mukhopadhyay, Nurul Nadzirin, Sreenath Nair, Typhaine Paysan-Lafosse, Lukas Pravda, David Sehnal, Osman Salih, Oliver Smart, James Tolchard, Mihaly Varadi, Radka Svobodova-Vařeková, Hossam Zaki, Gerard J Kleywegt, and Sameer Velankar. PDBe: improved findability of macromolecular structure data in the PDB. Nucleic Acids Research, 48(D1):D335–D343, January 2020. ISSN 0305-1048. doi: 10.1093/nar/gkz990. URL https://doi.org/10.1093/nar/gkz990.

David Baker and Andrej Sali. Protein structure prediction and structural genomics. Science, 294(5540):93–96, 2001. doi: 10.1126/science.1065659. URL https://www.science.org/doi/abs/10.1126/science.1065659.

Helen M. Berman, John Westbrook, Zukang Feng, Gary Gilliland, T. N. Bhat, Helge Weissig, Ilya N. Shindyalov, and Philip E. Bourne. The protein data bank. Nucleic Acids Research, 28 (1):235–242, 2000. ISSN 0305-1048. doi: 10.1093/nar/28.1.235. URL https://doi.org/10.1093/nar/28.1.235.

Graeme B. Bolger. The cAMP-signaling cancers: Clinically-divergent disorders with a common central pathway. Frontiers in Endocrinology, 13, October 2022. 1664–2392. doi: 10.3389/fendo.2022.1024423. https://www.frontiersin.org/journals/endocrinology/articles/10.3389/fendo.2022.1024423/full.

Carl Ivar Branden and John Tooze. Introduction to protein structure. Garland Science, 2012. ISBN 1136969888.

Stephen K Burley, Rusham Bhatt, Charmi Bhikadiya, Chunxiao Bi, Alison Biester, Pratyoy Biswas, Sebastian Bittrich, Santiago Blaumann, Ronald Brown, Henry Chao, Vivek Reddy Chithari, Paul A Craig, Gregg V Crichlow, Jose M Duarte, Shuchismita Dutta, Zukang Feng, Justin W Flatt, Sutapa Ghosh, David S Goodsell, Rachel Kramer Green, Vladimir Guranovic, Jeremy Henry, Brian P Hudson, Michael Joy, Jason T Kaelber, Igor Khokhriakov, Jhih-Siang Lai, Catherine L Lawson, Yuhe Liang, Douglas Myers-Turnbull, Ezra Peisach, Irina Persikova, Dennis W Piehl, Aditya Pingale, Yana Rose, Jared Sagendorf, Andrej Sali, Joan Segura, Monica Sekharan, Chenghua Shao, James Smith, Michael Trumbull, Brinda Vallat, Maria Voigt, Ben Webb, Shamara Whetstone, Amy Wu-Wu, Tongji Xing, Jasmine Y Young, Arthur Zalevsky, and Christine Zardecki. Updated resources for exploring experimentally-determined PDB structures and Computed Structure Models at the RCSB Protein Data Bank. Nucleic Acids Research, 53(D1):D564–D574, January 2025. ISSN 1362-4962. doi: 10.1093/nar/gkae1091. URL https://doi.org/10.1093/nar/gkae1091.

James J. Chou, Shipeng Li, Claude B. Klee, and Ad Bax. Solution structure of Ca2+–calmodulin reveals flexible hand-like properties of its domains. Nature Structural Biology, 8(11):990–997, November 2001. ISSN 1545-9985. doi: 10.1038/nsb1101-990. URL https://www.nature.com/articles/nsb1101-990.

P. J. Cock, T. Antao, J. T. Chang, B. A. Chapman, C. J. Cox, A. Dalke, I. Friedberg, T. Hamelryck, F. Kauff, B. Wilczynski, and M. J. de Hoon. Biopython: freely available python tools for computational molecular biology and bioinformatics. Bioinformatics, 25(11):1422–3, 2009. ISSN 1367-4811 (Electronic) 1367-4803 (Print) 1367-4803 (Linking). doi: 10.1093/bioinformatics/btp163. URL https://www.ncbi.nlm.nih.gov/pubmed/1930487

Cock, Peter J A Antao, Tiago Chang, Jeffrey T Chapman, Brad A Cox, Cymon J Dalke, Andrew Friedberg, Iddo Hamelryck, Thomas Kauff, Frank Wilczynski, Bartek de Hoon, Michiel J L eng R00 LM009837/LM/NLM NIH HHS/Research Support, Non-U.S. Gov’t England 2009/03/24 Bioinformatics. 2009 Jun 1;25(11):1422–3. doi: 10.1093/bioinformatics/btp163. Epub 2009 Mar 20.

Michel Ducreux, Ali Chamseddine, Pierre Laurent-Puig, Cristina Smolenschi, Antoine Hollebecque, Peggy Dartigues, Emmanuelle Samallin, Valérie Boige, David Malka, and Maximiliano Gelli. Molecular targeted therapy of BRAF-mutant colorectal cancer. Therapeutic Advances in Medical Oncology, 11:175883591985649, June 2019. doi: 10.1177/1758835919856494.

Douglas Hanahan. Hallmarks of cancer: New dimensions. Cancer Discovery, 12(1):31–46, 2022. ISSN 2159-8274. doi: 10.1158/2159-8290.Cd-21-1059. URL https://doi.org/10.1158/2159-8290.CD-21-1059.

John Jumper, Richard Evans, Alexander Pritzel, Tim Green, Michael Figurnov, Olaf Ronneberger, Kathryn Tunyasuvunakool, Russ Bates, Augustin Žídek, Anna Potapenko, Alex Bridgland, Clemens Meyer, Simon A. A. Kohl, Andrew J. Ballard, Andrew Cowie, Bernardino Romera-Paredes, Stanislav Nikolov, Rishub Jain, Jonas Adler, Trevor Back, Stig Petersen, David Reiman, Ellen Clancy, Michal Zielinski, Martin Steinegger, Michalina Pacholska, Tamas Berghammer, Sebastian Bodenstein, David Silver, Oriol Vinyals, Andrew W. Senior, Koray Kavukcuoglu, Pushmeet Kohli, and Demis Hassabis. Highly accurate protein structure prediction with alphafold. Nature, 596(7873):583–589, 2021. ISSN 1476-4687. doi: 10.1038/s41586-021-03819-2. URL https://doi.org/10.1038/s41586-021-03819-2.

A. Kryshtafovych, T. Schwede, M. Topf, K. Fidelis, and J. Moult. Critical assessment of methods of protein structure prediction (casp)-round xiii. Proteins, 87(12):1011–1020, 2019. ISSN 0887-3585 (Print) 0887-3585. doi: 10.1002/prot.25823.

A. Kryshtafovych, T. Schwede, M. Topf, K. Fidelis, and J. Moult. Critical assessment of methods of protein structure prediction (casp)-round xiv. Proteins, 89(12):1607–1617, 2021. ISSN 0887-3585 (Print) 0887-3585. doi: 10.1002/prot.26237.

Himansu Kumar, Lin-Ya Tang, Chengyuan Yang, and Pora Kim. Fusionpdb: a knowledgebase of human fusion proteins. Nucleic Acids Research, 52(D1):D1289–D1304, 2024. ISSN 0305-1048. doi: 10.1093/nar/gkad920. URL https://doi.org/10.1093/nar/gkad920.

Zeming Lin, Halil Akin, Roshan Rao, Brian Hie, Zhongkai Zhu, Wenting Lu, Nikita Smetanin, Robert Verkuil, Ori Kabeli, Yaniv Shmueli, Allan dos Santos Costa, Maryam Fazel-Zarandi, Tom Sercu, Salvatore Candido, and Alexander Rives. Evolutionary-scale prediction of atomic-level protein structure with a language model. Science, 379(6637):1123–1130, 2023. doi: 10.1126/science.ade2574. URL https://www.science.org/doi/abs/10.1126/science.ade2574.

Marc A. Martí-Renom, Ashley C. Stuart, András Fiser, Roberto Sánchez, Francisco Melo, and Andrej Šali. Comparative protein structure modeling of genes and genomes. Annual Review of Biophysics and Biomolecular Structure, 29(Volume 29, 2000):291–325, 2000. ISSN 1936-1238. doi: 10.1146/annurev.biophys.29.1.291. URL https://www.annualreviews.org/content/journals/10.1146/annurev.biophys.29.1.291.

Fredrik Mertens, Bertil Johansson, Thoas Fioretos, and Felix Mitelman. The emerging complexity of gene fusions in cancer. Nature Reviews Cancer, 15(6):371–381, 2015. ISSN 1474-1768. doi: 10.1038/nrc3947. URL https://doi.org/10.1038/nrc3947.

M. Mirdita, K. Schutze, Y. Moriwaki, L. Heo, S. Ovchinnikov, and M. Steinegger. Colabfold: making protein folding accessible to all. Nat Methods, 19(6):679–682, 2022. ISSN 1548-7105 (Electronic) 1548-7091 (Print) 1548-7091 (Linking). doi: 10.1038/s41592-022-01488-1. URL https://www.ncbi.nlm.nih.gov/pubmed/35637307.

Mirdita, Milot Schutze, Konstantin Moriwaki, Yoshitaka Heo, Lim Ovchinnikov, Sergey Steinegger, Martin eng DP5 OD026389/OD/NIH HHS/R21 AI156595/AI/NIAID NIH HHS/Research Support, N.I.H., Extramural Research Support, Non-U.S. Gov’t Research Support, U.S. Gov’t, Non-P.H.S. 2022/06/01 Nat Methods. 2022 Jun;19(6):679–682. doi: 10.1038/s41592-022-01488-1. Epub 2022 May 30.

Joannella Morales, Shashikant Pujar, Jane E. Loveland, Alex Astashyn, Ruth Bennett, Andrew Berry, Eric Cox, Claire Davidson, Olga Ermolaeva, Catherine M. Farrell, Reham Fatima, Laurent Gil, Tamara Goldfarb, Jose M. Gonzalez, Diana Haddad, Matthew Hardy, Toby Hunt, John Jackson, Vinita S. Joardar, Michael Kay, Vamsi K. Kodali, Kelly M. McGarvey, Aoife McMahon, Jonathan M. Mudge, Daniel N. Murphy, Michael R. Murphy, Bhanu Rajput, Sanjida H. Rangwala, Lillian D. Riddick, Françoise Thibaud-Nissen, Glen Threadgold, Anjana R. Vatsan, Craig Wallin, David Webb, Paul Flicek, Ewan Birney, Kim D. Pruitt, Adam Frankish, Fiona Cunningham, and Terence D. Murphy. A joint ncbi and embl-ebi transcript set for clinical genomics and research. Nature, 604(7905):310–315, 2022. ISSN 1476-4687. doi: 10.1038/s41586-022-04558-8. URL https://doi.org/10.1038/s41586-022-04558-8.

Saro Passaro, Gabriele Corso, Jeremy Wohlwend, Mateo Reveiz, Stephan Thaler, Vignesh Ram Somnath, Noah Getz, Tally Portnoi, Julien Roy, Hannes Stark, David Kwabi-Addo, Dominique Beaini, Tommi Jaakkola, and Regina Barzilay. Boltz-2: Towards accurate and efficient binding affinity prediction. bioRxiv, page 2025.06.14.659707, 2025. doi: 10.1101/2025.06.14.659707. URL https://www.biorxiv.org/content/biorxiv/early/2025/06/18/2025.06.14.659707.full.pdf.

Roshan M. Rao, Jason Liu, Robert Verkuil, Joshua Meier, John F. Canny, Pieter Abbeel, Tom Sercu, and Alexander Rives. Msa transformer, 2021. URL https://proceedings.mlr.press/v139/rao21a.html.

Alexander Rives, Joshua Meier, Tom Sercu, Siddharth Goyal, Zeming Lin, Jason Liu, Demi Guo, Myle Ott, C. Lawrence Zitnick, Jerry Ma, and Rob Fergus. Biological structure and function emerge from scaling unsupervised learning to 250 million protein sequences. Proceedings of the National Academy of Sciences, 118(15):e2016239118, 2021. doi: 10.1073/pnas.2016239118. URL https://www.pnas.org/doi/abs/10.1073/pnas.2016239118.

Carol A. Rohl, Charlie E. M. Strauss, Kira M. S. Misura, and David Baker. Protein Structure Prediction Using Rosetta, volume 383, page 66–93. Academic Press, 2004. ISBN 0076-6879. doi: 10.1016/S0076-6879(04)83004-0. URL https://www.sciencedirect.com/science/article/pii/S0076687904830040.

Arunachalam Sekar, Boobash Raj Selvadurai, Rishita Chatterjee, Lipika Pal, Aakanksha Verma, Nishanth Belugali Nataraj, Diana Drago Garcia, Suvendu Giri, Alessandro Genna, Feride Karatekin, Nitin Gupta, Deepthi Ramesh-Kumar, Mirie Zerbib, Yaron Vinik, Tamir Avioz, Eviatar Weizman, Eyal David, Alejandro A. Schäffer, Yunqian Pan, Haojie Huang, Wytske M. van Weerden, Eva Corey, Hazel Hunt, Andrew E. Greenstein, Ronnie Blecher-Gonen, Roni Oren, Ariel Afek, Ido Amit, Sima Lev, Anson Ku, Sumeyra Kartal, Jack R Bright, Rosina T Lis, William L Dahut, Adam G. Sowalsky, Eytan Ruppin, and Yosef Yarden. TMPRSS2-ERG confers resistance of prostate cancer to antiandrogens. EMBO Molecular Medicine, 18(6):2062–2097, June 2026. ISSN 1757-4684. doi: 10.1038/s44321-026-00423-7. URL https://doi.org/10.1038/s44321-026-00423-7.

UniProt Consortium. UniProt: the universal protein knowledgebase in 2025. Nucleic Acids Research, 53(D1):D609–D617, 2025. ISSN 0305-1048. doi: 10.1093/nar/gkae1010. URL https://doi.org/10.1093/nar/gkae1010.

Bert Vogelstein, Nickolas Papadopoulos, Victor E. Velculescu, Shibin Zhou, Luis A. Diaz, and Kenneth W. Kinzler. Cancer genome landscapes. Science, 339(6127):1546–1558, 2013. doi: 10.1126/science.1235122. URL https://www.science.org/doi/abs/10.1126/science.1235122.

Jeremy Wohlwend, Gabriele Corso, Saro Passaro, Noah Getz, Mateo Reveiz, Ken Leidal, Wojtek Swiderski, Liam Atkinson, Tally Portnoi, Itamar Chinn, Jacob Silterra, Tommi Jaakkola, and Regina Barzilay. Boltz-1: Democratizing biomolecular interaction modeling. bioRxiv, page 2024.11.19.624167, 2024. doi: 10.1101/2024.11.19.624167. URL https://www.biorxiv.org/content/biorxiv/early/2025/05/06/2024.11.19.624167.full.pdf.

P. E. Wright and H. J. Dyson. Intrinsically disordered proteins in cellular signalling and regulation. Nat Rev Mol Cell Biol, 16(1):18–29, 2015. ISSN 1471-0080 (Electronic) 1471-0072 (Print) 1471-0072 (Linking). doi: 10.1038/nrm3920. URL https://www.ncbi.nlm.nih.gov/pubmed/25531225.

Wright, Peter E Dyson, H Jane eng P01 GM071862/GM/NIGMS NIH HHS/R01 CA096865/CA/NCI NIH HHS/CA096865/CA/NCI NIH HHS/GM71862/GM/NIGMS NIH HHS/Research Support, N.I.H., Extramural Research Support, Non-U.S. Gov’t Review England 2014/12/23 Nat Rev Mol Cell Biol. 2015 Jan;16(1):18–29. doi: 10.1038/nrm3920.

Ruidong Wu, Fan Ding, Rui Wang, Rui Shen, Xiwen Zhang, Shitong Luo, Chenpeng Su, Zuofan Wu, Qi Xie, Bonnie Berger, Jianzhu Ma, and Jian Peng. High-resolution de novo structure prediction from primary sequence. bioRxiv, page 2022.07.21.500999, 2022. doi: 10.1101/2022.07.21.500999. URL https://www.biorxiv.org/content/biorxiv/early/2022/07/22/2022.07.21.500999.full.pdf.

Kurt Wüthrich. NMR with proteins and nucleic acids, volume 17 of Europhysics News. Wiley, New York, 1986. ISBN 0471828939.

J. Yang and Y. Zhang. Protein structure and function prediction using i-tasser. Curr Protoc Bioinformatics, 52:5.8.1–5.8.15, 2015. ISSN 1934-3396 (Print) 1934-3396. doi: 10.1002/0471250953.bi0508s52.

Juwon Yoon, Sang-Hyeok Lee, Soohan Moon, Harerta Gebremedhn, Teklab Gebregiworgis, and Ki-Young Lee. Integrative structural and dynamics studies of epidermal growth factor receptor (EGFR). Experimental & Molecular Medicine, 58(6):1711–1720, Jun. 2026. ISSN 2092-6413. doi: 10.1038/s12276-026-01744-w. URL https://www.nature.com/articles/s12276-026-01744-w.

Chengxin Zhang, Morgan Shine, Anna Marie Pyle, and Yang Zhang. Us-align: universal structure alignments of proteins, nucleic acids, and macromolecular complexes. Nature Methods, 19(9):1109–1115, 2022. ISSN 1548-7105. doi: 10.1038/s41592-022-01585-1. URL https://doi.org/10.1038/s41592-022-01585-1.

Xiyu Zhao, Victor B. Yang, Arjun K. Menta, Jacob Blum, Adam Wahida, and Vivek Subbiah. Alphafold’s predictive revolution in precision oncology. AI in Precision Oncology, 1(3):160–167, 2024. doi: 10.1089/aipo.2024.0010. URL https://journals.sagepub.com/doi/abs/10.1089/aipo.2024.0010.

